# OpenCRS: an open-source regulated human cardiorespiratory model with large-scale calibration and global sensitivity analysis at rest and during exercise

**DOI:** 10.64898/2026.09.14.751454

**Authors:** Sheng-Ya Wang, Harry Saxton, Maximilian Balmus, Steven A. Niederer

## Abstract

Cardiopulmonary exercise testing reveals cardiovascular and respiratory limitations not apparent at rest, but similar measurements can arise from different interacting regulatory mechanisms, preventing physiological causal inference from data alone. Mechanistic computational models can separate these mechanisms *in silico*. However, whole-body cardiorespiratory models contain hundreds of parameters, making global sensitivity analysis and calibration challenging. We present OpenCRS, an open-source Python cardiorespiratory model coupling closed-loop 0D lumped-parameter circulation with gas exchange and integrated autonomic and respiratory control. The framework incorporates baroreflex, chemoreflex, pulmonary stretch receptors, central command, and neuromuscular drive, together with novel representations of atrial dynamics and exercise baroreflex set-point resetting. We introduce a scalable calibration pipeline that (i) applies a derivative-based global sensitivity measure (DGSM) directly to the simulator, reducing 272 parameters to 72 influential ones; (ii) trains Gaussian process emulator surrogates within iterative History Matching to exclude implausible parameter regions; and (iii) performs Bayesian calibration (MCMC), inferring a joint posterior with Hamiltonian Monte Carlo (No-U-Turn sampler) under a Gaussian copula prior. A single maximum a posteriori parameter set simultaneously reproduced 50 literature-derived rest and exercise targets (45/50 within 1 SD, all within 1.83 SD), with the rest-to-exercise transition emerging from the model’s embedded feedback rather than independent fitting. Simulator-based DGSM agreed with constrained Sobol indices (mean Spearman rank correlation 0.79). Sensitivity analysis identified influential physiological mechanisms. Only 2–16 parameters contributed *>*2% of the Sobol total-effect sensitivity per target. Resting cardiovascular targets were driven by unstressed volumes and cardiac mechanics, while exercise shifted influence towards autonomic efferent regulation. Respiratory outputs remained most sensitive to chemoreflex and gas exchange parameters. These shifts capture coordinated cardiorespiratory adaptation to metabolic demand. More broadly, the framework provides a population-level prior with quantified uncertainty for cardiovascular digital twins and a reusable route for calibrating high-dimensional, regulated physiological models.

**Author summary:** Exercise reveals limits of the heart and lungs that resting measurements can miss, but the same result can arise from several interacting processes. We built OpenCRS, a computer model linking the heart, blood circulation, lungs, and the body’s automatic control of breathing and blood pressure, to separate these possible causes. Such models contain hundreds of adjustable parameters, making it challenging to identify which most influence measurements and which combinations fit the data. We developed an efficient workflow that screened a 272 parameter model to retain 72 influential ones, ruled out combinations that could not produce realistic results, and estimated uncertainty in the remaining values. We then identified a set of values that best reproduced 50 measurements at rest and during exercise. We found that resting heart function was shaped mainly by blood volume and how the heart fills and pumps, while nervous-system control becomes more influential during exercise. Breathing and gas exchange remained strongly affected by the body’s responses to oxygen and carbon dioxide. Our model and workflow provide a reusable basis for testing explanations of changes in heart and lung function and developing personalised computer models to study health and disease.

## Introduction

Clinical decisions are often informed by resting hospital measurements, but it is during exercise that the dynamically regulated cardiorespiratory system can reveal limitations in performance and functional capacity. Exercise assessments are used for disease diagnosis and prognosis, with testing modalities exposing physiological abnormalities at the early stages of pathological adaptation [1]. Cardiopulmonary exercise testing (CPET) can infer reduced contractile reserve and impaired lusitropy, which are early markers of right ventricular dysfunction in pulmonary arterial hypertension, while in heart failure with preserved ejection fraction, CPET-measured VO_2_max provides prognostic value by indicating limitations across the entire oxygen transport pathway [1,2]. However, isolating the coupled mechanisms shaping the observed data for therapeutic targeting remains challenging. The same clinical measurement can arise from various underlying processes, such as homeostatic control compensating for disturbances in one subsystem by adjusting another, or noisy data with limited resolution masking meaningful differences [3]. During exercise, increased metabolic demand also shifts the relative influence of subsystems on measurements so that different regulatory mechanisms dominate under stress [4]. This motivates the identification of key drivers in observed metrics across rest and exercise to support targeted intervention. Computational mathematical models address this challenge, assimilating mechanistic relationships from physiology, physics, and anatomy into a quantitative framework [5].

Such integrative models enable patient-specific simulations that evaluate cardiorespiratory function and test competing mechanistic explanations *in silico*, recognising pathophysiological insights obscured in observable measurements alone, and informing both personalised treatment strategies and generalisable trends [5]. An extensive cardiorespiratory model integrates a multi-scale framework of tissue and fluid mechanics, ventilation dynamics, electrophysiological ion channel behaviour, and subcellular chemical signalling pathways to capture cardiac and pulmonary function [6]. Prior cardiovascular modelling studies have spanned single or multi-scale interactions coupled across various dimensions [7, 8]. 3D models resolve local flow and spatial geometry to quantify metrics such as wall shear stress that is associated with endothelial dysfunction and vascular remodelling [9, 10]. 2D models simulate planar cell deformation, multiphase flows, and flow-induced shear stress, while 1D models consider pressure and flow-wave propagations in only the axial direction to examine global haemodynamics [10]. Although computationally efficient relative to 3D models, they remain too resource-intensive for real-time clinical decision-making [6]. 0D lumped parameter models (LPMs) have gained traction in addressing this limitation by averaging out spatial variation to compartmentalise and simulate bulk flow, pressure, and volume across cardiac chambers and the circulatory system. The cardiovascular system is represented as an electric circuit. Flow rate is analogous to current, blood pressure to voltage, hydraulic resistances account for compartment pressure losses, and compliance describes the blood volume stored at a given pressure [6]. LPMs support investigations of pathology evolution, the evaluation of physiological or surgical impacts, and are boundary conditions for higher-dimensional models [11–14].

LPMs are systems of ordinary differential equations (ODEs), in which biomarker rate-of-change equations are integrated over time from initial conditions to generate dynamic and continuous traces [6]. Cardiovascular LPMs simulate cardiac function over successive cycles, capturing physiological adaptations such as exercise, standing, or sleep [15–17]. The ODEs are derived from mass conservation through the continuity equation, and Poiseuille-based momentum conservation, relating blood flow to pressure gradients [18].

Whole human circulation LPMs embody the cardiovascular system as interconnected compartments. The Ursino & Magosso (1998-2002) models include the systemic and pulmonary circulations given by Windkessel models [19–21]. The four heart chambers are simulated by a time-varying elastance model introduced by Suga et al. (1974), postulating that the contractile state (elastance) is represented by a load-independent time-varying elastance function, defining the instantaneous pressure-volume (PV) relationship for a given inotropic state [22]. Ursino & Magosso models also consider baroreflex regulation, driving adaptations in peripheral resistance, unstressed volumes (blood volume not contributing to flow), heart rate, and elastances [21]. Although widely used for cardiovascular control, these models rely largely on reflex parameters derived from canine experiments, leaving human values uncertain [21]. Subsequent models have the same Windkessel circulation foundation but focus on different aspects of the cardiovascular system. Elstad et al. (2002) modelled exercise baroreflex resetting and D’Angelo and Papelier (2005) modelled the chemoreflex regulation of tissue blood flow [23, 24]. Later, Shi and Korakianitis (2006) improved valve dynamics, and CircAdapt by Lumens et al. (2009) incorporates left-right ventricular wall interactions [25, 26]. Fernandes et al. (2024) introduced a detailed baroreflex mechanism using the Hodgkin-Huxley model for afferent and central neurons [27]. While these cardiovascular models were appropriate for resting and haemodynamic applications, omitting respiratory mechanics overlooks the respiratory control of cardiovascular dynamics during exercise [28].

Current cardiorespiratory models by Cheng et al. (2010), Albanese et al. (2016), Serna et al. (2018), and Carlos et al. (2022) couple respiratory mechanics, respiratory control, and gas exchange with Windkessel cardiovascular models [15, 17, 29, 30]. PNEUMA (Cheng et al. (2010)) incorporates autonomic regulation of peripheral resistances, unstressed volumes, heart rate, and elastances by baroreceptors, chemoreceptors, pulmonary stretch receptors, and the CNS response to arterial pCO_2_ and pO_2_ [15]. Its sleep mechanisms also recognise circadian and ultradian rhythms. Sarmiento et al. (2021) adapted PNEUMA for exercise by adding breathing-work minimisation to optimise breathing profiles, according to gas-regulated alveolar flow (refer to Section 2.1.4) [17]. It showed better agreement with experimental exercise data than the Albanese et al. (2016) model, which assumed a fixed inspiratory-to-expiratory ratio and was designed for hypercapnic and hypoxic stimuli [17]. The Sarmiento model also had lower cardiovascular prediction errors than the respiratory-focused Serna et al. (2018) model and is proprietary, with no open-source availability [17]. Across these models, parameter identification remained challenging because many parameters have no clinically interpretable surrogates, existing to fit experimental curves rather than being directly measurable. Clinical data availability is therefore essential for physiological parameter calibration.

Sensitivity analyses provide a structured approach for identifying the parameters and subsystems that most influence measured metrics, supporting clinicians in determining therapeutic targets to prioritise. Local SAs are generally derivative-based, assessing how small perturbations of each parameter around its nominal value affect the output while holding others constant [31]. Parameter interactions are ignored, and failure to consider the full input parameter range prevents the model from characterising the complete output variability. This contrasts with global sensitivity analyses (GSA), which explore the entire input parameter space. Sobol GSA, a variance-based method, is the predominant technique [31]. The output variance is decomposed into first-order effects, the direct influence of a parameter on the output variance, and higher-order effects, accounting for parameter-parameter interactions. A Sobol GSA requires *N* (*P* + 2) simulations, where *P* is the number of parameters, and *N*, the number of parameter samples, can be over 100,000 to reach a converged periodic state, as reported by Tlalka et al. (2024) for a one chamber regulated model with 36 parameters. This is computationally challenging for models with over 200 parameters [4].

The Derivative-based GSA measure (DGSM) is an efficient global approach for screening non-influential parameters [32]. DGSM provides an upper bound on the Sobol total-order effect 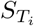 for an output *y* and parameter *θ*_*i*_, without explicitly decomposing sensitivities into first-order *S*_*i*_, second-order *S*_*ij*_, and higher-order interaction effects (Eq. (1)) [33]. By avoiding the full variance decomposition, DGSM can screen a high-dimensional parameter space with fewer model evaluations. Reducing the parameter set used to capture output metrics allows the model to retain mechanistic complexity while focusing tuning on a smaller subset of more identifiable, physiologically meaningful parameters.

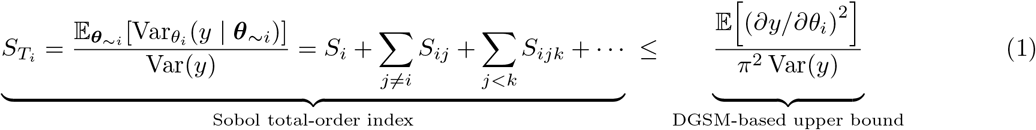

The first objective of the current work is to present OpenCRS, a novel feedback-regulated cardiorespiratory model, the simulator, that integrates established submodels with physiologically grounded augmentations. The 272-parameter simulator couples cardiovascular circulation, gas exchange, and cardiorespiratory controllers into a unified framework with improved atrial, breathing, and contraction mechanics. Exercise is stimulated by increasing metabolic O_2_ consumption and CO_2_ production, with exercise-specific adaptations in ventilatory and autonomic control that modulate contractility, heart rate, unstressed volumes, resistances, and ventilation. This shifts resting regulation toward stressed-state physiology.

The second objective is to perform DGSM directly on the simulator as a fast first-pass parameter screening tool over the full, global parameter space with all 272 parameters varied by ±50% of their nominal values, to reduce dimensionality before calibration. Previous GSAs have used Gaussian Process emulators (GPEs) to approximate the simulator input-output relationships [34, 35]. However, it is computationally impractical to train a reliable GPE in 272 dimensions. GPEs also impose local smoothness and continuity, returning outputs even for parameter sets that cause simulator failures, discontinuities, or instability [36]. DGSM instead derives sensitivities by efficiently identifying a subset of influential parameters from the simulator’s true input-output mapping. For verification, DGSM was compared against Sobol GSA over the reduced 72 influential parameters, further constrained to the physiologically non implausible region.

The third objective is to simultaneously calibrate the model against 50 representative literature-derived targets measured at rest and during exercise. The calibration workflow does not treat rest and exercise as independent fitting problems; instead, it tests whether parameter sets consistent with resting physiology can reproduce the adaptation to exercise and its targets via the model’s embedded physiological feedback and control mechanisms. As such, the framework demonstrates physiological consistency across states.

Calibrated models with quantified parameter and predictive uncertainty are a prerequisite for cardiovascular digital twins, which are repeatedly updated with longitudinal data to track and predict patient-specific physiology [37]. At scale, these twins could be updated sequentially as new data arrive, rather than recalibrated from scratch [38]. This requires an informative starting point, which our joint, uncertainty-aware calibration provides as a population-level prior for individual updating.

## Materials and methods

We first describe OpenCRS, the feedback-regulated cardiorespiratory simulator, in Section 2.1 (Fig 1), followed by the calibration pipeline in Section 2.2 (Fig 2). Calibration aimed to characterise the joint distribution of parameter sets consistent with 50 physiological targets at rest and during exercise, and to identify the single best-fitting parameter set. DGSMs were computed for these targets using simulator evaluations that continued directly from convergence at rest to convergence following exercise perturbation. The resulting sensitivities identified a reduced subset of influential parameters (Section 2.2.1). History Matching then used GPEs to efficiently search within the reduced parameter space and iteratively rule out implausible regions given the rest and exercise outputs (Section 2.2.2). MCMC subsequently used GPEs trained within the physiologically plausible space to sample the remaining Not-Ruled-Out-Yet (NROY) region and identify the best-fitted parameter set, defined by the highest joint posterior density. Finally, Sobol GSA quantified parameter influence and compared sensitivity patterns within this NROY region.

**Fig 1.**
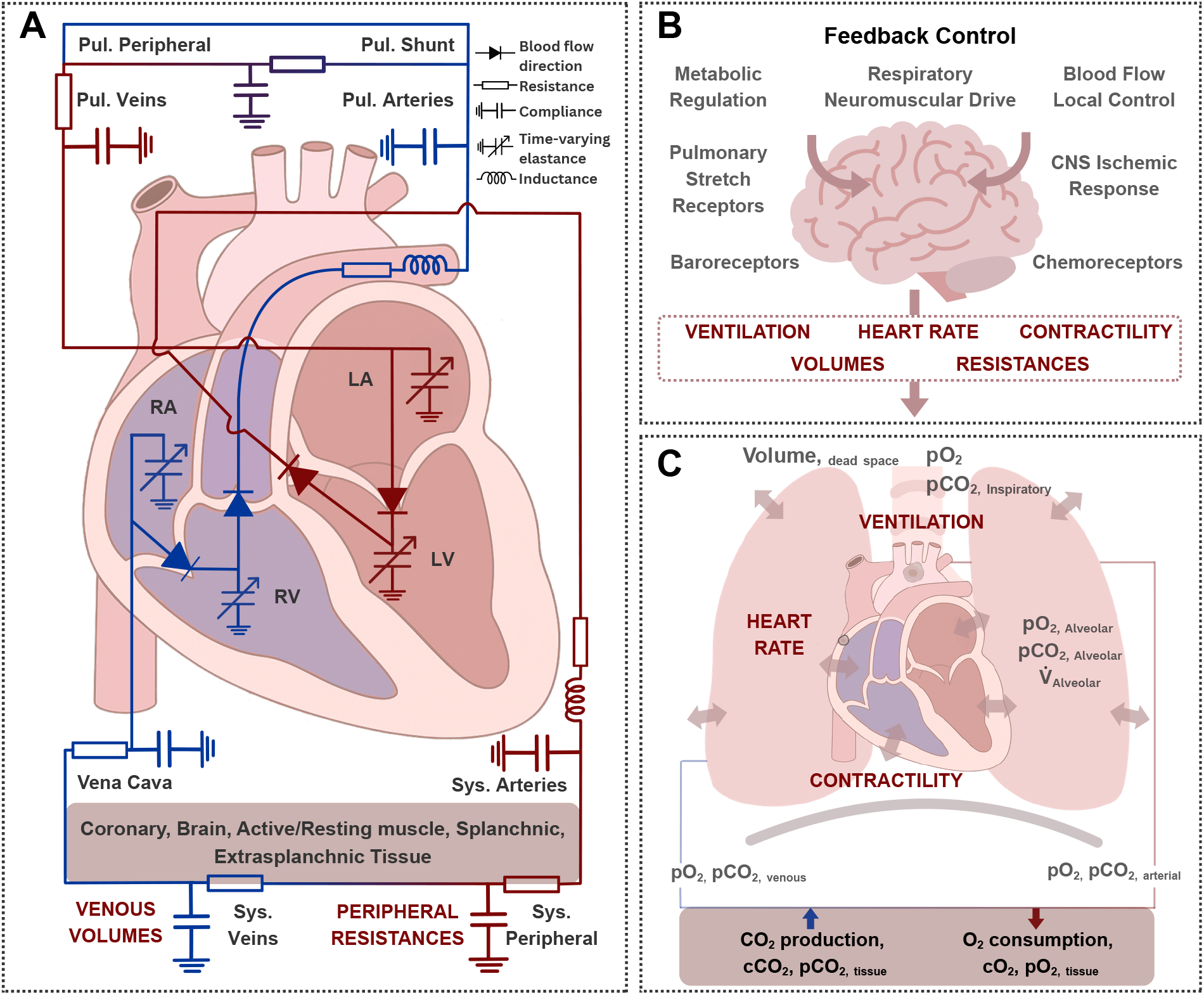
OpenCRS schematic. A: Cardiovascular LPM with the systemic and pulmonary circulation. B: Feedback mechanisms modulate ventilation, heart rate, contractility, volumes, and resistances. C: Pulmonary LPM with blood gas concentrations linked to alveolar ventilatory partial pressures.

**Fig 2.**
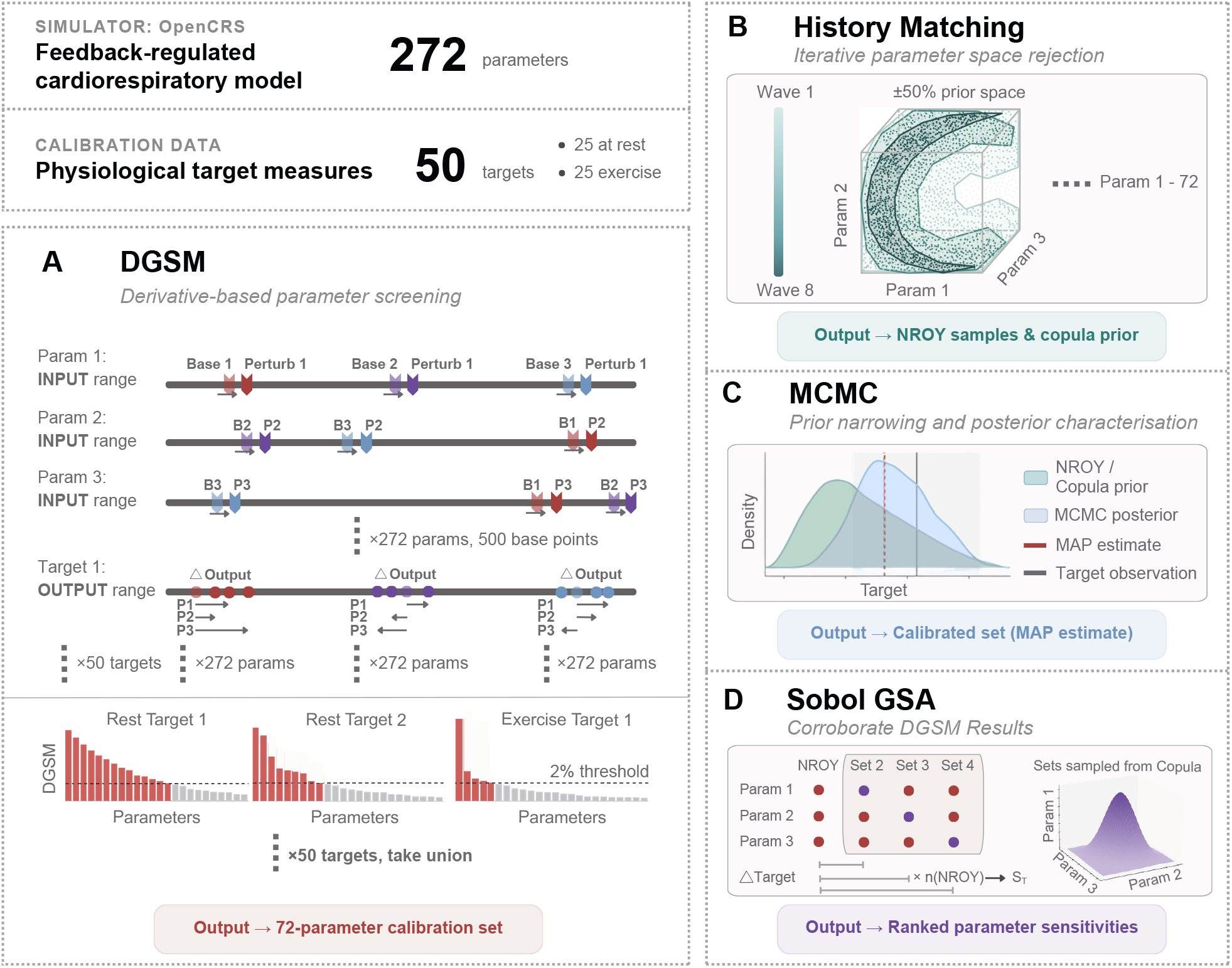
Calibration workflow. A: Derivative-based global sensitivity analysis (DGSM) screens the 272-parameter simulator, retaining the union of parameters contributing ≥2% of the total DGSM for any of the 50 targets (25 at rest, 25 with exercise). B: History Matching trains Gaussian-process emulators and progressively excludes implausible parameter regions, narrowing the Not-Ruled-Out-Yet (NROY) space over successive waves. C: MCMC using the No-U-Turn Sampler (NUTS) samples the joint posterior under a copula prior fitted to the NROY region, with the likelihood measuring agreement between emulator predictions and calibration targets. The maximum a posteriori (MAP) estimate is taken as the best-calibrated parameter set. D: Sobol total-order (*S*_*T*_) and first-order (*S*_*i*_) effects are compared with DGSM to investigate influential parameter shifts and mechanistic drivers of observed metrics.

### 2.1 Cardiorespiratory model

The cardiorespiratory model unifies literature submodels and adds physiologically grounded augmentations into a novel open-source Python framework. The cardiovascular system and controller follow Ursino & Magosso (2002) with parameter values taken from Sarmiento et al. (2021) [17, 19]. The implemented pressure-volume, contraction, and valvular flow model was described by Suga and Sagawa (1974) and Korakianitis and Shi (2005) [22, 39]. Novel modelling approaches for atrial dynamics and resetting the baroreceptor set point are described. The gas exchange models combine formulations from Sarmiento et al. (2021) and Chiari et al. (1997), while the respiratory controller introduces new adaptations to the Serna et al. (2017) approach [17, 40, 41]. All 272 parameter values, definitions, original literature values, and parameter adjustments with justifications are described in S1 Appendix. Manual adjustments to literature parameter values were minimal and explicitly documented to demonstrate a systematic and reproducible approach to model calibration.

#### 2.1.1 The cardiovascular model

The cardiovascular model (Fig 1A) represents the systemic and pulmonary circulations as Windkessel compartments [19]. Blood is ejected from the heart into the systemic arteries, passes through the peripheral and venous compartments, and returns via the vena cava to enter the pulmonary circulation. Systemic circulation is divided into six compartments: coronary, brain, resting muscle, active muscle, splanchnic, and extrasplanchnic vascular beds, each with its own peripheral and venous compartments, allowing distinct control mechanisms to be assigned to each (S1 Appendix, Section 1).

For each model compartment (*i*), state variables define pressures, flows, and volumes. Volume equations follow mass conservation, with the inflow (*Q*_*i*−1_) minus outflow (*Q*_*i*_) characterising the change in volume (*dV/dt*) (Eq (2)). Compliance (*C*_*i*_) relates pressure (*P*_*i*_) changes to volume changes (Eq (3)). The flow, pressure, and resistance (*R*_*i*_) relationship is analogous to Ohm’s law (Eq (4)), and the systemic and pulmonary arteries includes inertance (*L*_*i*_) for a four-element Windkessel formulation (Eq (5)) [17]. Compartment pressures adopt a linear pressure-volume relationship with constant compliance until they fall below external pressure. At this point, vessels act as Starling resistors, and flow is limited by the external tissue pressure. This increases effective venous resistance, leading to the venous waterfall phenomenon (S1 Appendix, Section 1.3) [42].

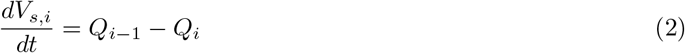

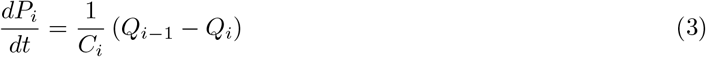

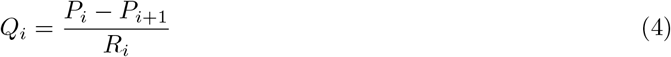

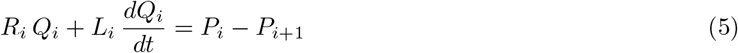

The time-varying elastance model determines instantaneous pressures of the ventricle and atrial chambers as a weighted combination of the end-systolic and end-diastolic pressure-volume relationships (ESPVR and EDPVR) (Fig 3A, Eq (6)) [6]. The ESPVR represents the active pressure contribution, modelled as a linear function of chamber elastance and volume relative *V*_0_, whereas the EDPVR is the passive pressure contribution modelled as an exponential relation for chamber filling, with parameters *c* and *K* fitted to the observed data (Eq (6)) [6]. The weighting between these active and passive components is governed by a double-cosine activation function that describes the systolic time course within each heartbeat (*t*_*i*_) (Eq (7)) [39].

**Fig 3.**
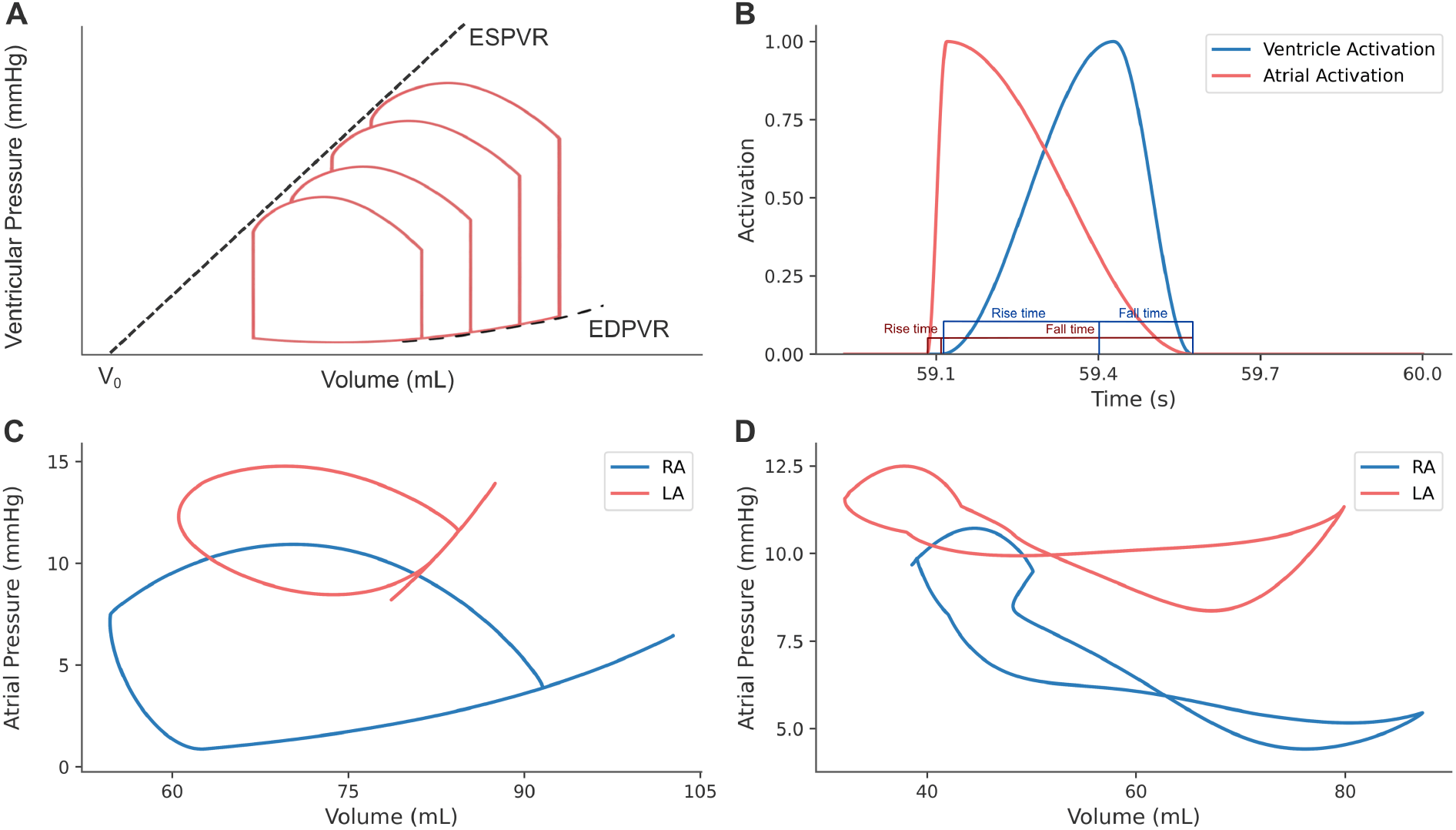
Dynamic cardiac chamber mechanics. A: Ventricular pressure-volume curves following the linear end-systolic and exponential end-diastolic pressure-volume relationship (ESPVR, EDPVR). B: Atrial and ventricular activation with adjustable rise and fall times. C: Atrial PV loop without a V wave, and D: with a V wave, after introducing pericardial effects and prolonging atrial relaxation in the activation function.

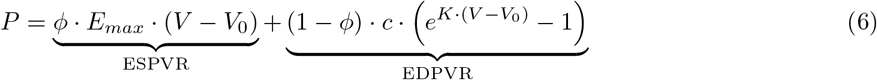

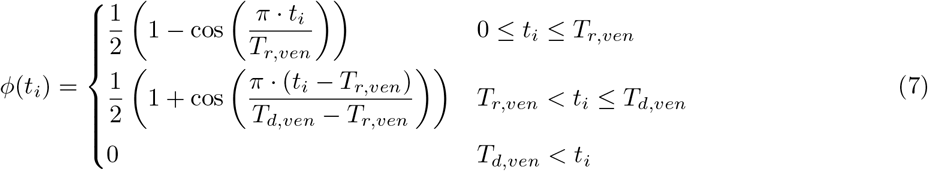

*T*_*r,ven*_ and *T*_*d,ven*_ denote systolic rise and fall times, with *ϕ* = 0 during diastole. Atrial activation is time shifted so atrial contraction occurs before ventricular systole. This contraction formulation has the advantage of adjustable delays, rise, and fall times (Fig 3B) [39]. The Korakianitis and Shi orifice model was adopted for physiological blood flow through the valves (S1 Appendix, Section 1.9) [26].

In atrial PV relationships, previous studies have ignored the V wave during atrial diastole by using the same PV relationship for both the conduit and reservoir phases (Fig 3C) [6, 43]. Models that reproduce the atrial figure-of-eight loop do so phenomenologically, linking elastance to hysteresis and piecewise volume-triggered changes rather than the physiological V-wave drivers of ventricular contraction and atrioventricular interaction [44, 45]. OpenCRS introduces pericardial pressure into the instantaneous PV relationship (Eq (8)) to account for atrioventricular coupling, and extends the atrial fall time in the atrial activation function. The pericardium mechanically constrains cardiac filling to prevent unrealistic chamber overexpansion, particularly the excessive atrial expansion just before atrial contraction [46]. Pericardial pressure is modelled here as a function of the heart chamber and pericardial volumes, according to an exponential PV relationship, with *V*_*offset*_ defining the cardiac volume at which pressure rises, and *V*_*scale*_ the steepness of the increase [47]. Elongation of the atrial fall time reflects the slower decay of the calcium transient and sarcomere-generated tension during the atrial reservoir phase [48]. This sustained activation is absent in the conduit phase, producing distinct conduit vs. reservoir dynamics, and the V-wave loop (Fig 3D).

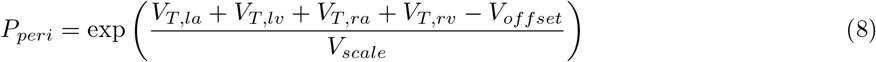

#### 2.1.2 The respiratory model

The current respiratory system combines Sarmiento et al. (2021) for airflow between the mouth and the alveoli, with Chiari et al. (1997) for lung, tissue, and brain gas exchange, and metabolic O_2_ and CO_2_ turnover [17, 40] (Fig 1C). The cardiovascular and respiratory model is coupled via the interaction between blood O_2_ and CO_2_ concentrations and their partial pressures (pO_2_/pCO_2_), with parameters fitted to follow literature O_2_ and CO_2_ dissociation curves (S1 Appendix, Section 4.1) [40, 49].

Gases move through conducting airways, modelled as a series of dead-space compartments, and partial pressure changes are proportional to the airflow and pressure gradients between compartments. During inspiration, O_2_-rich gas propagates towards the alveoli, while during expiration, CO_2_-rich alveolar gas cascades back to the mouth, capturing convective delay and sequential mixing (Fig 4) [17].

**Fig 4.**
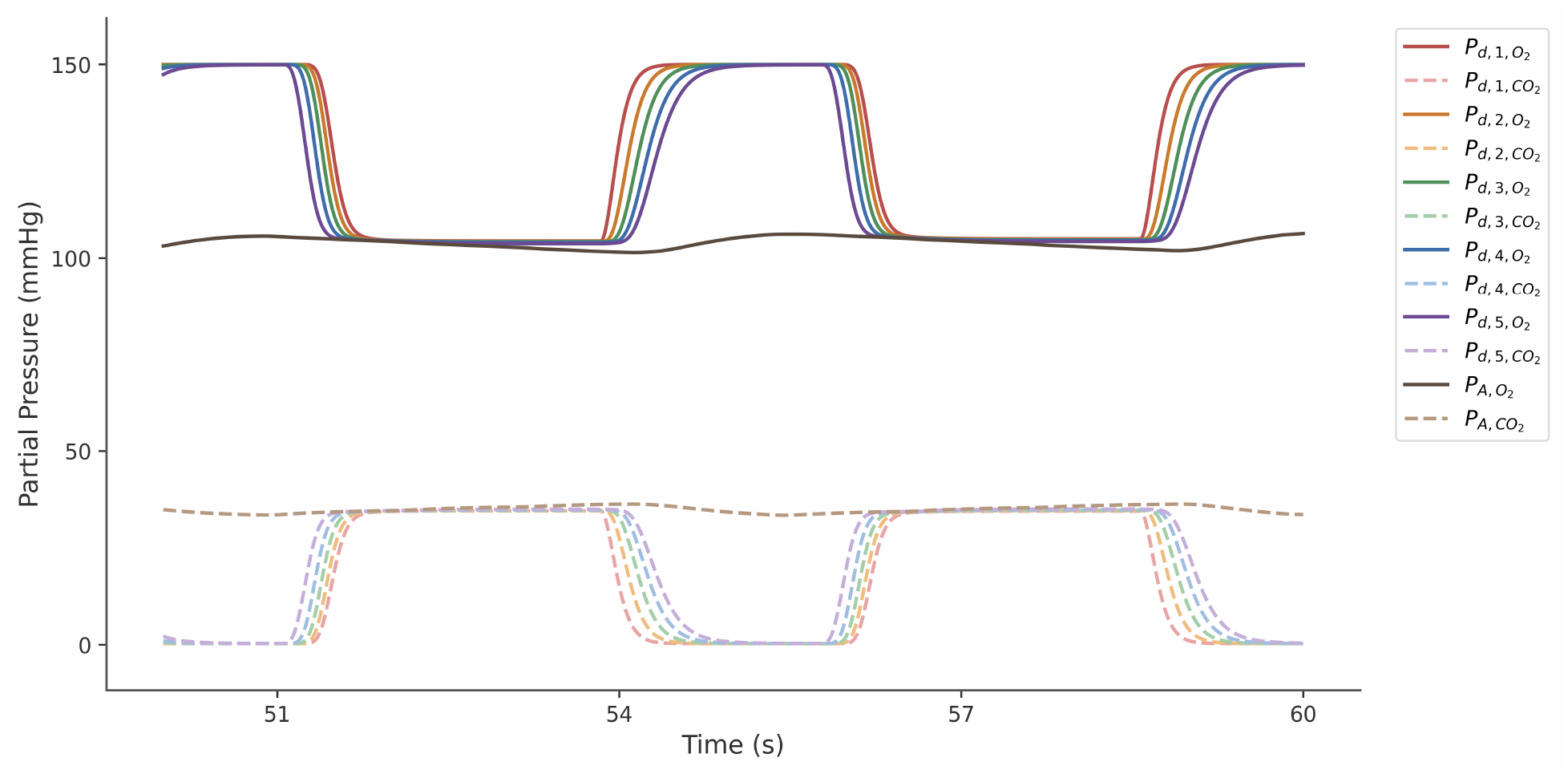
Oxygen and carbon dioxide partial pressures in the conducting airways. Partial pressures are shown for the conducting airway compartments *P*_*d*1_ to *P*_*d*5_ and for the alveolar gas partial pressures (PAO_2_ and PACO_2_) across three breaths.

At the alveoli, alveolar-capillary gas transfer is governed by perfusion and ventilation dynamics. This establishes a mass-balance formulation for cardiovascular-to-respiratory coupling. Perfusion is represented by Fick’s principle, where gas transfer is proportional to pulmonary peripheral blood flow (*Q*) and the capillary-arterial blood concentration gradient (*C*_*v,gas*_ −*C*_*a,gas*_), while ventilation accounts for bulk convective transport between the alveolar compartment (*P*_*A,gas*_) and the conducting airway compartments (*P*_*d*(*X*)*gas*_) (Eq (9)). The alveolar ventilation equation (Eq (10)) uses the factor 863 mmHg to convert the perfusion term’s volumetric blood gas content to a pressure-based form and from standard to body gas conditions consistent with Eq (9) (S1 Appendix, Section 4.1) [50]. This captures the inflow of inspired gas to the alveoli during inspiration and the return of alveolar gas to the airways during expiration [50, 51].

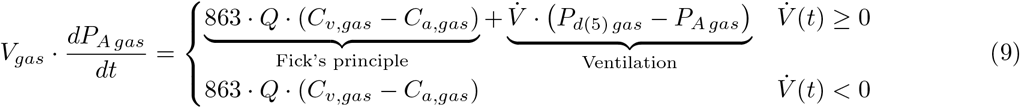

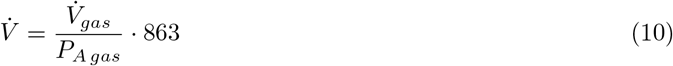

OpenCRS implements the Chiari et al. (1997) gas exchange model (Fig 5). This explicitly accounts for O_2_ dissolved in plasma versus bound to hemoglobin, the pulmonary shunt for blood bypassing gas exchange, and enforces mass balance (Eq (11)) across distinct brain and tissue compartments, each with its own arterial and venous circulations (S1 Appendix, Section 4.2) [40].

**Fig 5.**
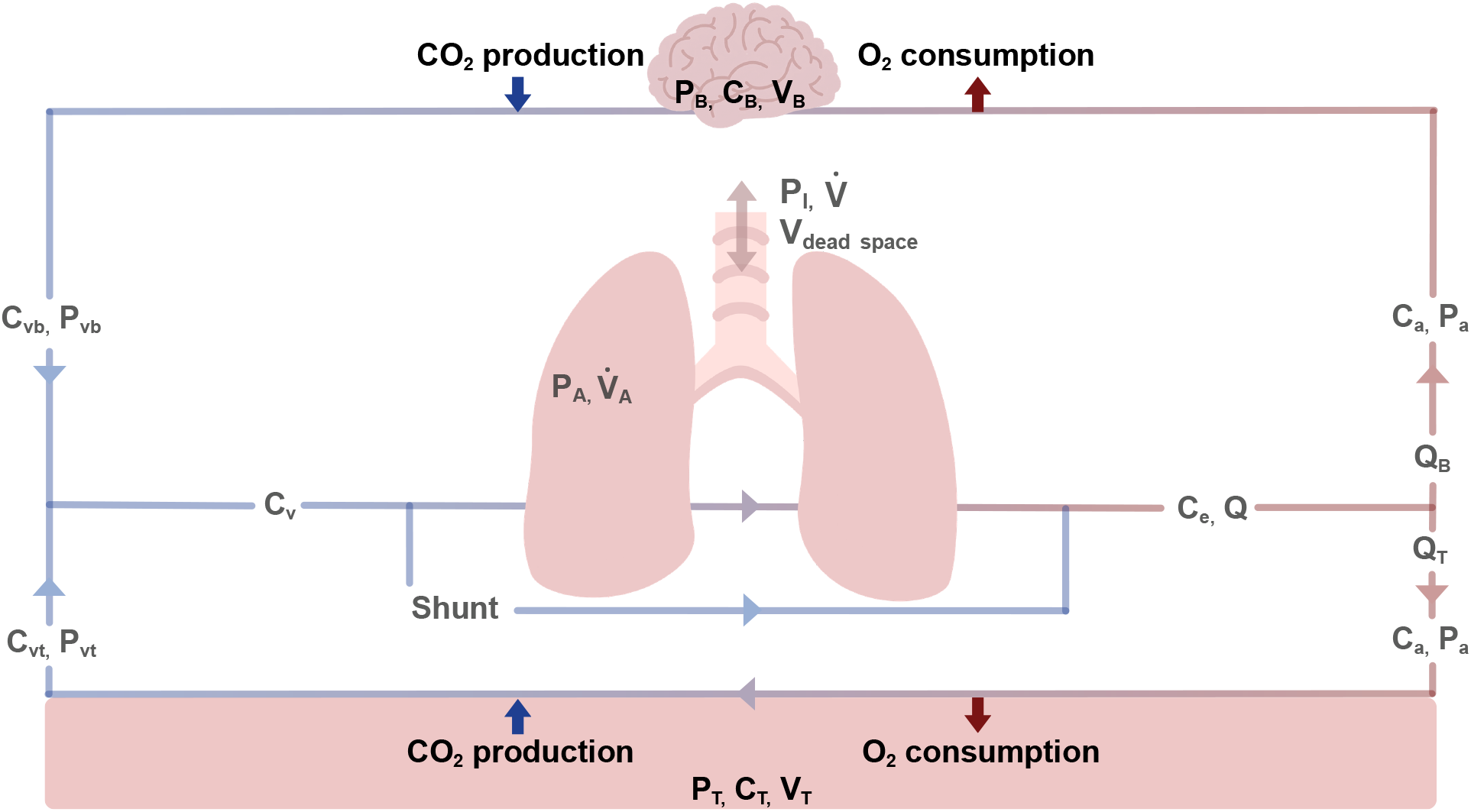
Pulmonary model with ventilation and perfusion coupling between the lungs, brain, and tissue. P: partial pressure, C: concentration, V: volume, 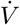: ventilation, Q: flow, I: inspiration, A: alveolar, B: brain, T: tissue, v: venous, a: arterial, e: lung exit.

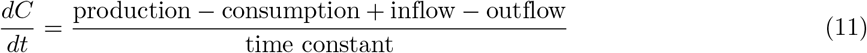

#### 2.1.3 The cardiovascular controller

Exercise is initiated by raising tissue metabolic CO_2_ production and O_2_ consumption rates to 80% of the anaerobic threshold (*AT*), simulating moderate-intensity exercise. Feedback mechanisms modelled by Magosso and Ursino (2002) and Sarmiento et al. (2021) allow the model to adapt dynamically to exercise [19]. Mechanisms include autonomically regulated pathways, local control of blood flow, central nervous system ischemic response, and respiratory neuromuscular drive, which together modulate ventilation, heart rate, contractility, resistance, and venous blood flow volumes (Fig 1B, Table 1).

**Table 1.**
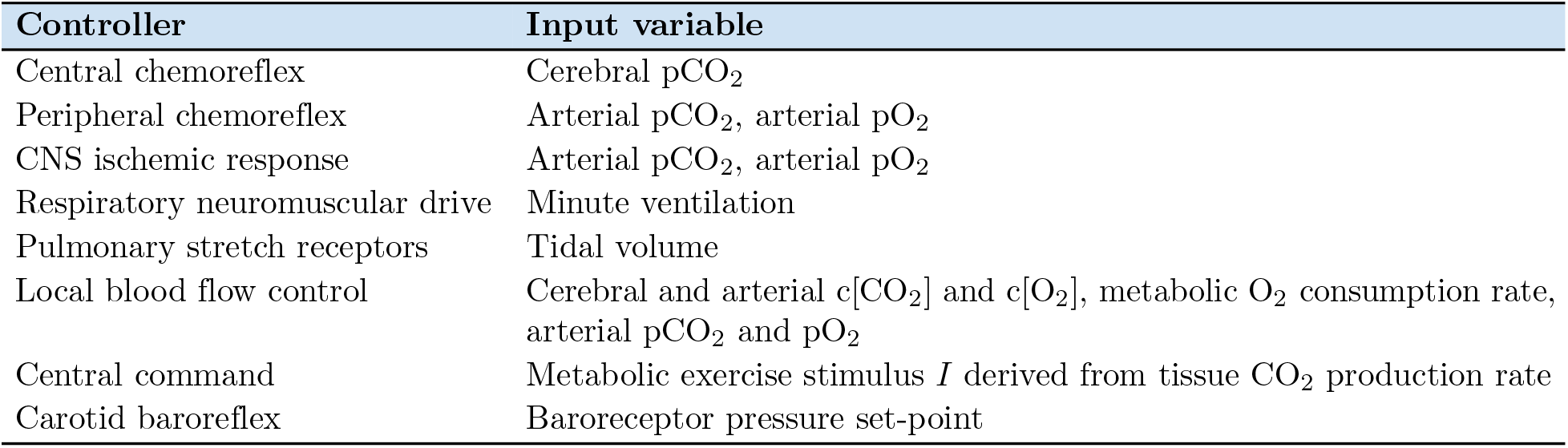
Feedback pathways and their determinants of firing rate.

| Controller | Input variable |
| --- | --- |
| Central chemoreflex | Cerebral $\text{pCO}_2$ |
| Peripheral chemoreflex | Arterial $\text{pCO}_2$, arterial $\text{pO}_2$ |
| CNS ischemic response | Arterial $\text{pCO}_2$, arterial $\text{pO}_2$ |
| Respiratory neuromuscular drive | Minute ventilation |
| Pulmonary stretch receptors | Tidal volume |
| Local blood flow control | Cerebral and arterial $\text{c}[\text{CO}_2]$ and $\text{c}[\text{O}_2]$, metabolic $\text{O}_2$ consumption rate, arterial $\text{pCO}_2$ and $\text{pO}_2$ |
| Central command | Metabolic exercise stimulus $I$ derived from tissue $\text{CO}_2$ production rate |
| Carotid baroreflex | Baroreceptor pressure set-point |

As metabolic activity rises, accumulating cerebral and arterial CO_2_ and falling O_2_ concentrations intensify central and peripheral chemoreceptor firing, increasing ventilation [52]. Peripheral chemoreceptor activation also increases both sympathetic and parasympathetic activity to promote vasoconstriction and bradycardia [52]. In addition, severe hypoxia and hypercapnia trigger the CNS ischemic response for a marked increase in sympathetic activity [17]. The respiratory neuromuscular drive increases in parallel with ventilatory demand, shaping the respiration-linked autonomic modulation associated with respiratory sinus arrhythmia, while pulmonary stretch receptors mediate the Hering-Breuer reflex to limit lung overinflation [53, 54]. Local blood flow control is achieved through vasomotor regulation of vascular resistance, with arterial oxygen and carbon dioxide levels driving adjustments in vascular tone to redistribute perfusion toward metabolically active tissues [17]. Concurrently, sympathetic venoconstriction shifts unstressed venous volumes into the circulation, raising venous return and cardiac output. Increasing tissue CO_2_ production rates (MRTCO_2_) with exercise reinforces the integrated cardiorespiratory response, in which central command directly modulates autonomic outflow. The dimensionless metabolic exercise intensity stimulus (*I*), calculated from the current MRTCO_2_, drives the central command and ranges from 0 at rest to 1 at *AT* (Eq (12)) [19].

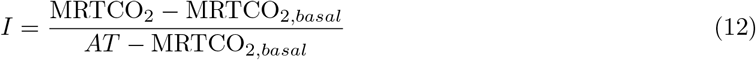

At the onset of exercise, the baroreceptor set point increases to shift the pressure-firing curve to a higher operating point. This prevents the higher pressures reached during exercise from evoking the usual baroreflex-mediated vagal activation and the sympathetic inhibition that would otherwise slow the heart and reduce blood pressure, allowing sympathetic drive to remain elevated. To recognise this mechanism, the target *P*_*n*_ varies with *I* between the nominal and maximum set point (Eq (13)), physiologically linking exercise intensity and central command to progressive baroreflex resetting [55]. The current set point approaches this target through a first-order low-pass filter with time constant *τ*_*p*_ (Eq (14)).

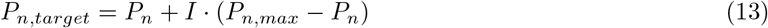

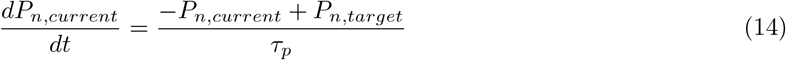

All feedback mechanisms share the same underlying framework, a monotonic sigmoidal function with a low-pass first-order filter determining the receptor firing rate (S1 Appendix, Section 2) [17]. The low-pass filter smooths the firing response with a time constant, conveying the gradual adjustment of firing towards the target value. The target firing is a sigmoid function with a maximum, minimum, and centre point. For each controller in Table 1, firing rates are combined into a weighted efferent signal that regulates peripheral resistances, venous volumes, heart rates, and contractilities. The feedback updates once per heartbeat based on the average efferent firing over that cycle [17].

#### 2.1.4 The respiratory controller

The respiratory controller adjusts breathing patterns by estimating alveolar ventilation from the mean gas partial pressures per breath [56]. From the alveolar and dead-space flows, the subsequent breath is determined by minimising the work of breathing, consistent with Serna et al. (2017) [41]. This procedure optimises airflow profiles and specifies inspiratory and expiratory times, tidal volume, and dead space volume (Fig 6).

**Fig 6.**
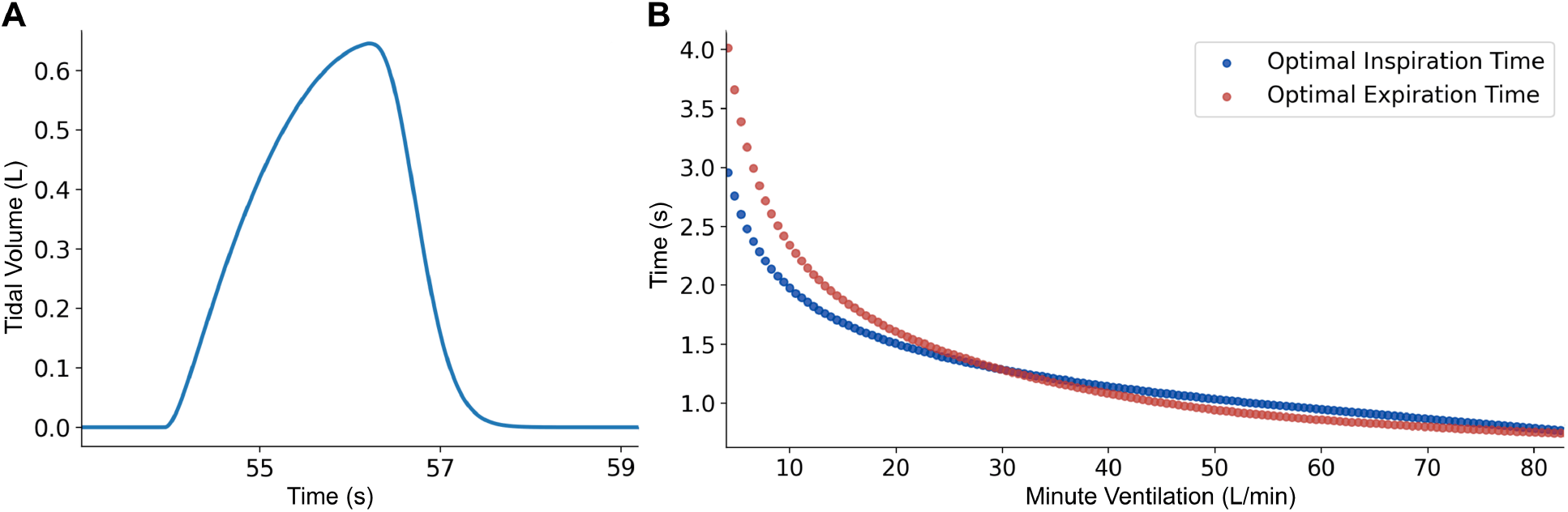
Optimised breathing pattern. A: Tidal volume. B: Nelder-Mead optimised breath timings.

The model extends Serna et al. (2017) with several augmentations. Respiratory muscle pressure remains quadratic during inspiration, but is modelled with a Gaussian rather than a negative exponential function during expiration. This enforces C1 continuity at the inspiration-expiration boundary and reduces spiking in the second derivative, improving numerical smoothness for integration and optimisation. Formally, we have

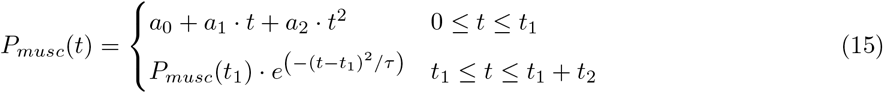

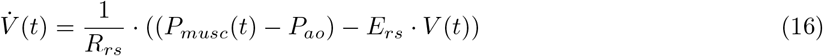

Imposing continuity at the inspiration-expiration boundary and setting *V* (*t*) to the tidal volume at end-inspiration yields closed-form solutions for the coefficients *a*_1_ and *a*_2_, respectively, and taking *τ* to approach zero at end-expiration reduces the optimisation to only the inspiration (*t*_1_) and expiration (*t*_2_) times (S1 Appendix, Section 3). Optimisation used the Nelder-Mead simplex method, a derivative-free method for piecewise objective functions [57]. Nelder-Mead produced more reproducible solutions across initialisations, and a smoother, more reliable convergence behaviour. To avoid performing an optimisation every breath, at the start of the simulation, optimal *t*_1_ and *t*_2_ were precomputed over the physiological range of minute ventilation, and a cubic spline fit provided a fast lookup mapping from minute ventilation to breathing times.

#### 2.1.5 Solver and performance time

The model was implemented in Python 3.12 with 82 state variables (S2 Appendix) and 272 parameters (S1 Appendix), and was solved using the adaptive explicit Runge-Kutta Bogacki-Shampine 3(2) (RK23) method. The maximum integration step was 1 ms, with relative and absolute tolerances of 1 *×* 10^−3^ and 1 *×*10^−6^, respectively. For each target, the simulator returned the mean (*µ*(*θ*)) over the final 10 cardiac cycles. Across History Matching, each parameter set was simulated at rest for 450 s, and limit cycle convergence was defined as a heart-rate range *<*2 bpm across the final 10 beats. If unmet, simulations were extended in 60-s increments up to 690 s. Exercise was then initiated and simulated for 700 s. The same convergence criterion was applied, with 60-s extensions up to 940 s after exercise onset. Total simulation time ranged from 1150 to 1630 s, depending on time to limit cycle convergence.

To improve performance, delayed feedback and moving-average values from the controllers were stored in preallocated arrays indexed by a shared dictionary. Each functioned as a circular buffer overwriting the oldest values once the buffer limit was reached, avoiding dynamic memory allocation. Delayed quantities required at the current iteration were retrieved by linear interpolation between the nearest stored times, providing a continuous lookup. The circular buffer was rewound when the solver rejected trial steps.

To further reduce computational cost, the repeatedly evaluated derivative functions were just-in-time (JIT) compiled using Numba v0.61.2’s @njit decorator. [58]. JIT compilation converts supported Python functions to optimised machine code at runtime. Subsequent calls are executed directly on the compiled code, unlike standard interpreted Python, where the interpreter executes bytecode line by line with additional overhead [58]. Because SciPy’s `solve ivp()` executes its step loop in interpreted Python, the RK23 step loop was also JIT-compiled from a transcription of SciPy’s implementation, reproducing its trajectories while returning only the final state. In a single-threaded benchmark, a 1000-s rest–exercise simulation required 27 s with JIT compilation versus 757 s without, running approximately 37 times faster than real time.

### 2.2 Calibration

Sensitivity analyses and calibration were interpreted with respect to 50 targets for rest and moderate-intensity exercise in a representative 70 kg adult male, with literature reference ranges (Table 2). Indexed quantities, such as volumes normalised to body surface area, were converted to absolute values for this reference individual (BSA: 1.73 m^2^, age: 25) [59]. Male-specific values were used to align with this reference case; however, many exercise studies did not report sex-stratified or indexed data. Therefore, the final reference set combines direct literature values, converted indexed values, and non-sex-stratified data. Other clinical measures including cardiac output (HR(EDV-ESV)), pericardial volume change (Δ*V*_RA+LA+RV+LV_), stroke volume (EDV-ESV), and ejection fraction ((EDV-ESV)/EDV), were excluded from calibration because they are derived from primary targets and accurate calibration of the underlying outputs constrains these quantities.

**Table 2.** Literature ranges for target clinical measures at rest and during exercise.

| Target | Rest (± SD) | Source | Exercise (± SD) | Source |
| --- | --- | --- | --- | --- |
| Heart Rate (HR) | 74 ± 13 bpm | [60] | 155 ± 21 bpm | [60] |
| LV Systolic Pressure (LV SP) | 123 ± 18 mmHg | [60] | 165 ± 23 mmHg | [60] |
| LV Diastolic Pressure (LV DP) | 76.7 ± 8.1 mmHg | [61] | 76.4 ± 9.1 mmHg | [61] |
| LV End-Systolic Volume (LV ESV) | 62.3 ± 15.6 mL | [62] | 45.5 ± 8.7 mL | [62] |
| LV End-Diastolic Volume (LV EDV) | 152.1 ± 27.7 mL | [62] | 145.5 ± 26.1 mL | [62] |
| RV Systolic Pressure (RV SP) | 22.5 ± 7.5 mmHg | [63] | 29.5 ± 7.5 mmHg* | [64] |
| RV Diastolic Pressure (RV DP) | 4 ± 3 mmHg | [63] | 9.9 ± 5.6 mmHg | [65] |
| RV End-Diastolic Volume (RV EDV) | 151.9 ± 31.7 mL | [62] | 139.4 ± 26.1 mL | [62] |
| RV End-Systolic Volume (RV ESV) | 64.4 ± 17.3 mL | [62] | 40.3 ± 10.6 mL | [62] |
| Max RA Pressure A wave (RA P<sub>max, A</sub>) | 8 ± 3 mmHg | [66] | 12 ± 4 mmHg | [66] |
| Max RA Pressure V wave (RA P<sub>max, V</sub>) | 5 ± 3 mmHg | [66] | 11 ± 4 mmHg | [66] |
| Max RA Volume (RA V<sub>max</sub>) | 92.4 ± 19.5 mL | [62] | 77.3 ± 18.5 mL | [62] |
| Min RA Volume (RA V<sub>min</sub>) | 45.7 ± 11.2 mL | [62] | 27.9 ± 5.0 mL | [62] |
| RA Pre-Atrial Contraction Vol. (RA V<sub>pre-A</sub>) | 57.4 ± 9.8 mL** | [62, 67] | 40.3 ± 6.0 mL** | [62, 68] |
| Max LA Pressure A wave (LA P<sub>max, A</sub>) | 13 ± 3 mmHg*** | [69] | 19 ± 7 mmHg*** | [69] |
| Max LA Pressure V wave (LA P<sub>max, V</sub>) | 12 ± 3 mmHg*** | [69] | 19 ± 8 mmHg*** | [69] |
| Max LA Volume (LA V<sub>max</sub>) | 68.3 ± 17.5 mL | [62] | 66.3 ± 19.7 mL | [62] |
| Min LA Volume (LA V<sub>min</sub>) | 30.6 ± 9.2 mL | [62] | 23.0 ± 9.7 mL | [62] |
| LA Pre-Atrial Contraction Vol. (LA V<sub>pre-A</sub>) | 40.0 ± 8.2 mL** | [62, 67] | 33.8 ± 8.8 mL** | [62, 68] |
| Max LV Pressure Derivative (LV dP/dt) | 1461 ± 383 mmHg/s | [70] | 1750 ± 522 mmHg/s | [70] |
| Max RV Pressure Derivative (RV dP/dt) | 271 ± 55 mmHg/s | [71] | 713 ± 110 mmHg/s | [71] |
| Tidal Volume (V<sub>T</sub>) | 850 ± 400 mL | [72] | 2220 ± 640 mL | [73] |
| Minute Ventilation (V<sub>E</sub>) | 11.4 ± 3.9 L/min | [73] | 62.6 ± 17.9 L/min | [73] |
| PaO<sub>2</sub> | 102.3 ± 11.2 mmHg | [74] | 97.2 ± 6.0 mmHg | [74] |
| PaCO<sub>2</sub> | 35.5 ± 4.9 mmHg | [74] | 38.4 ± 2.6 mmHg | [74] |

\* Increased the rest value by 7 mmHg, matching the average exercise-induced rise in RV SP [64]. \*\* V<sub>pre-A</sub> was derived from $V_{pre-A} = V_{min} + f(V_{max} - V_{min})$ , assuming atrial contraction contributes 20%–30% to ventricular filling ( $f$ ), and SDs estimated by error propagation. Because literature V<sub>pre-A</sub> is inconsistently defined and tightly coupled to V<sub>max</sub> and V<sub>min</sub>, it was not a target for calibration. Instead, V<sub>pre-A</sub>, V<sub>max</sub>, and V<sub>min</sub> emulators estimate $f$ , with values within the physiological 20%–30% range considered acceptable [67]. In History Matching, uncertainty in $f$ was propagated using Monte Carlo draws from the emulator predictive distributions and samples retained if $P(0.20 \leq f \leq 0.30) \geq 0.05$ . In MCMC, $f$ was centered at 0.25, and the V<sub>pre-A</sub>, V<sub>min</sub>, and V<sub>max</sub> joint emulator uncertainty for $f$ was propagated into the likelihood using deterministic cubature. \*\*\* Pulmonary arterial wedge pressures were taken as LA pressure surrogates [69].

#### 2.2.1 Derivative-based global sensitivity measure (DGSM)

DGSM quantifies, on average, how sensitive a target output is to small perturbations in each parameter across the input space. The model was evaluated at 500 base parameters sets (base points, *y*(*θ*)), with each parameters perturbed about every base point (*y*(*θ* + Δ*θ*)) [32]. Each base point was first simulated following Section 2.1.5, but using a 300-s rest period. Its converged resting state then initialised all runs in that block, including the unperturbed run, providing *y*(*θ*) and *y*(*θ* + Δ*θ*) with a common warm start. Each run re-converged at rest for a further 300 s, recorded resting targets, and simulated 700 s of exercise, with 60-s extensions up to 940 s. DGSM (*ν*_*j*_) values were computed from finite differences gradient estimates (*dy/dθ*) (Eq (17)), the lower (*a*_*j*_) and upper (*b*_*j*_) bounds of parameter *j*, and output variance (Var(*y*(*θ*))) across the 500 base-point outputs (Eq (18)). Averaging yielded the mean DGSM, and bootstrapping estimated the variability. Larger DGSM values indicated more influential parameters whose perturbations produce larger changes in the target output. Parameters contributing at least 2% of total DGSM were retained; the remainder screened out as having negligible influence over the specified range.

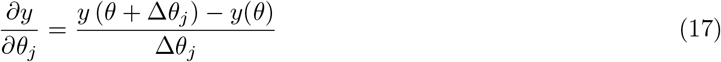

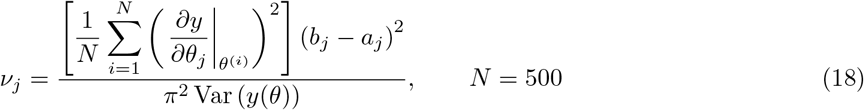

To set up the DGSM pipeline, each model parameter was bounded at ±50% around its nominal value, defining the hypercube [*a*_*j*_, *b*_*j*_] used for sampling. Base samples were generated using the finite difference sampler from SALib v1.5.1. This draws *N* quasi-random base points and, for each base sample, perturbs the parameters individually (Δ*θ*), producing (1 + no. of parameters) simulations per base sample and ((1 + no. of parameters) · no. of base samples) total samples. We modified SALib to scale bootstrap uncertainty to the DGSM and to exclude finite-difference pairs with atypical step sizes or squared-derivative contributions (*>*3 SD), which otherwise amplified numerical artefacts (S3 Appendix, Section 1).

While 500 base parameter sets were used as a conservative choice, to determine the minimum number of base points required for consistently retained parameters, DGSM estimates were evaluated using increasing numbers of base points at intervals of 10. Convergence was defined as the earliest base point after which no new parameters were found to add to the cumulative union, with consecutive Jaccard similarity used as an additional measure of parameter-set stability.

To assess robustness to parameter ranges and ranking choices, the DGSM analysis was repeated over parameter hypercubes, [*a*_*j*_, *b*_*j*_], spanning 20% and 50% below and above their nominal values. Consistent influential parameters with widening ranges were interpreted as evidence that the screening results were not overly dependent on chosen bounds. Conversely, substantial changes indicate model nonlinearity and range dependence, as well as uncertainty-driven effects, where parameters assigned wider uncertainty ranges are expected to generate greater output variability and may therefore appear more influential.

#### 2.2.2 History matching

The History Matching (HM) workflow is an iterative process in which successive “waves”rule out implausible regions of the parameter space inconsistent with observations. Parameters contributing at least 2% of the total output DGSM for each target were retained. The final calibration parameter set was then defined as the union of retained parameters across all 50 target outputs (Table 2). Parameters outside this subset were fixed to their nominal (midpoint) values, while retained parameters were assigned uniform prior bounds of ±50%.

The full parameter space was sampled using emulators for each target observation as low-cost surrogates of the full model. The Autoemulate v1.1.1 Python package was used to identify the Gaussian process emulator with a constant mean function and Matérn covariance kernel of smoothness 1.5 as the emulator (GPE) with the most accurate predictive performance [75]. Initial GPEs were trained on 8192 simulations using a Latin Hypercube (LHC) experimental design, with all parameters varied by ±50% of their nominal values.

In the first wave, 400,000 LHC samples were drawn from the ±50% uniform prior and evaluated using the GPEs to broadly explore the calibration parameter space. For each parameter set, emulator predictions were compared with physiological target values using an implausibility measure:

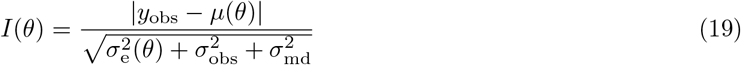

Where *y*_obs_ is the physiological target value, *µ*(*θ*) the emulator prediction at parameter set *θ*, 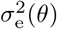 the emulator predictive variance, and 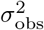 the observation variance. The model discrepancy term 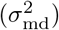 was set to zero, assuming the simulator captured the dominant physiology for the target outputs considered, and any residual structural error is small relative to the observation variance. A parameter set was retained and classified as NROY if the maximum implausibility across all emulator-predicted outputs did not exceed 3.25 in the first wave and a threshold of 3 in the following waves, as per Pukelsheim’s 3*σ* rule [34]. Additional filters rejected emulator-predicted samples with nonphysiological atrial volumes, ensuring atrial contraction contributed 20%–30% to ventricular filling, positive minimum resting atrial volumes, and exercise atrial volumes above 10 mL, as the 3*σ* bounds otherwise permitted negative volumes [67]. 6000 randomly selected NROY parameter sets were then evaluated by the simulator, and remaining valid simulations after applying the above atrial filters were used to retrain emulators for the next wave.

In subsequent waves, 200,000 parameter sets were generated for emulator evaluation via cloud sampling. For each NROY parameter set, a truncated multivariate normal distribution was constructed with its mean vector equal to that parameter set. The standard deviation for each parameter was set to a fixed fraction (0.1) of its range in the current NROY sample. Truncation prevented proposed samples from falling outside the prior parameter bounds. Equal numbers of new parameter sets were drawn independently around each NROY parameter set to form the 200,000 parameter sets. As in the first wave, each following wave simulated 6000 random NROY parameter sets and refit the emulators on the resulting valid simulations. This targets exploration within the surviving plausible region, with the NROY space progressively narrowing across successive waves. HM was terminated at the wave in which the median emulator predictive uncertainty across all outputs was below 10% of the observation uncertainty.

#### 2.2.3 MCMC

In a patient-specific setting, the objective is to infer parameter values that best reconcile the model with an individual’s measured data. HM delineates a non-implausible region for the parameter space, and Markov chain Monte Carlo (MCMC) completes calibration by characterising the joint posterior distribution over the parameters conditioned on the targets. The joint posterior quantifies the probability distribution of parameter combinations consistent with the target data, and its joint mode, the maximum a posteriori estimate (MAP), provides the single best-supported parameter set.

For a proposed parameter set *θ* and target outputs ***y***_obs_, the posterior distribution was defined:

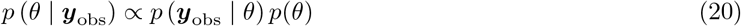

Where *p* (***y***_obs_ | *θ*) is the likelihood and *p*(*θ*) is the prior density. To quantify agreement between the emulator predictions and the *n*_targets_ = 50 targets, a Gaussian joint likelihood was adopted:

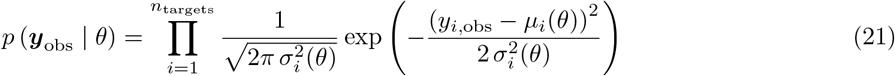

With total variance:

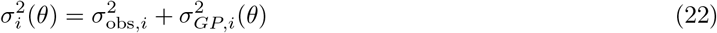

Where 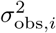 is the target variance for output *i*, and *µ*_*i*_(*θ*) and 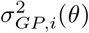 are the predictive mean and variance of the corresponding Gaussian process emulator from the final wave of HM. Sampling was carried out on the log-posterior scale, so the product likelihood reduced to a sum of per-target log-likelihood contributions. This Gaussian likelihood served as a pragmatic calibration criterion that assigns greater posterior support to parameter sets whose predicted outputs are closer to the targets, while simultaneously accounting for both measurement and emulator uncertainty.

The prior density was modelled using a Gaussian copula fitted to the discrete NROY samples, representing the joint distribution as univariate marginal distributions and a dependence structure [76]. Each parameter’s marginal distribution was modelled using a bounded log-spline density, and the samples were transformed to a Gaussianised coordinate system to represent joint dependence by a multivariate normal and correlation matrix [77]. This construction preserves the non-Gaussian marginal shapes of the NROY region while also encoding the correlation structure.

Posterior samples were drawn using the No-U-Turn Sampler (NUTS) as implemented in Pyro v1.9.1. NUTS exploits gradients of the log-posterior to explore the high-dimensional parameter space more efficiently than random-walk samplers [78]. It extends Hamiltonian Monte Carlo by adaptively determining the trajectory length at each iteration, removing the need to pre-specify the number of leapfrog steps [78]. Because the log-posterior is a differentiable function of the parameters through the GPE, prior, and likelihood, the gradients were obtained by automatic differentiation. Four chains were initialised at a random NROY parameter sample to ensure sampling began within a non-implausible region. To allow sampling in an unbounded coordinate space, each bounded parameter was transformed using a logit mapping before running NUTS. Chains were run with 500 warm-up iterations followed by 3000 posterior samples per chain, and convergence and sampling efficiency were assessed for each calibration parameter using Pyro’s Gelman-Rubin statistic (split 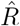), and Geyer’s autocorrelation-based effective sample size (ESS) [79, 80].

After MCMC, the posterior draw with the highest log posterior across all four chains was selected as the initial maximum a posteriori (MAP) estimate and refined using local gradient-based L-BFGS-B optimisation. The optimised parameter values were taken as the best-calibrated set, while non-calibrated parameters retained their original nominal values.

#### 2.2.4 Sobol GSA

To corroborate the DGSM method, we perform Sobol GSA to compare parameter sensitivities between rest and exercise with correlated parameter inputs within the physiologically plausible target space. The analysis used the final NROY parameter sets from HM (63721), defined by an implausibility threshold below 3. Conventional Saltelli column swapping is inappropriate for this analysis because independently replacing one parameter while holding the remaining fixed destroys the dependence structure induced by the physiological constraints. We therefore adapted the constrained Sobol GSA of Kucherenko et al. (2017). For each target (*Y*), the dependent-input first-order effect (*S*_*i*_) used:

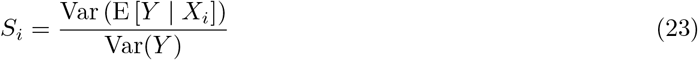

With E [*Y* | *X*_*i*_] approximated by ordering the NROY samples by the magnitude of parameter *X*_*i*_ and averaging emulator predictions within equally populated bins [81]. *S*_*i*_ ranks parameters by how strongly *X*_*i*_ varies the output. Because the remaining parameters are not fixed, it includes information carried through parameter dependence. Following Kucherenko et al. (2017) and setting roughly as many bins as observations, *S*_*i*_ was estimated using 225 equally populated bins, containing ~225 NROY points each [81].

The total-order effect (*S*_*T*_) used:

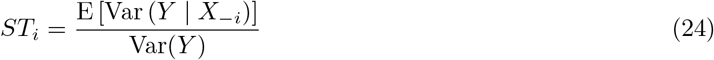

*S*_*T*_ was calculated with each NROY point as an anchor. For parameter *X*_*i*_, the remaining parameters *X*_−*i*_ were held constant while another 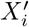 value was drawn from the conditional distribution *X*_*i*_ *X*_−*i*_, implied by a Gaussian copula with logspline marginals fitted to the NROY sample and treated as the dependent joint distribution of the model inputs [81–83]. Parameter sets were reassessed against the implausibility threshold with the emulator and redrawn when outside the NROY region. This formed paired parameter sets at every NROY anchor and *S*_*T*_ was estimated as the mean squared output difference over 2Var(*Y*) [81]. Var(*Y*) was calculated across all NROY parameter sets for each target. This conditional resampling approximates the distribution within the feasible, non-rectangular NROY domain, preserving the structural dependencies induced by the physiological output constraints.

## Results

The following sections present the results of the calibration framework, progressing from DGSM through HM and MCMC to obtain a final calibrated parameter set consistent with both rest and exercise targets. Sobol sensitivity analysis, evaluated over plausible parameter sets, was used as a complementary variance-based method to support the sensitivity patterns identified by DGSM.

### 3.1 DGSM

DGSM results are shown for the ±50% parameter perturbation range, with the ±20% and base-point convergence analyses provided in S3 Appendix, Sections 2 and 3, respectively. DGSM reduced the model’s effective dimensionality by identifying a subset of 72 out of 272 influential parameters across the 50 reported rest and exercise targets. For each target, parameters that contributed at least 2% of the total DGSM were retained. Fig 7 summarises these parameter-output relationships, highlighting the parameters dominating the cardiorespiratory target outputs. Despite the high-dimensional parameter space, all target outputs were dominated by only a few parameters. S3 Appendix, Section 4 reports exact DGSM values, cumulative DGSM coverage, and 95% bootstrap confidence intervals.

**Fig 7.**
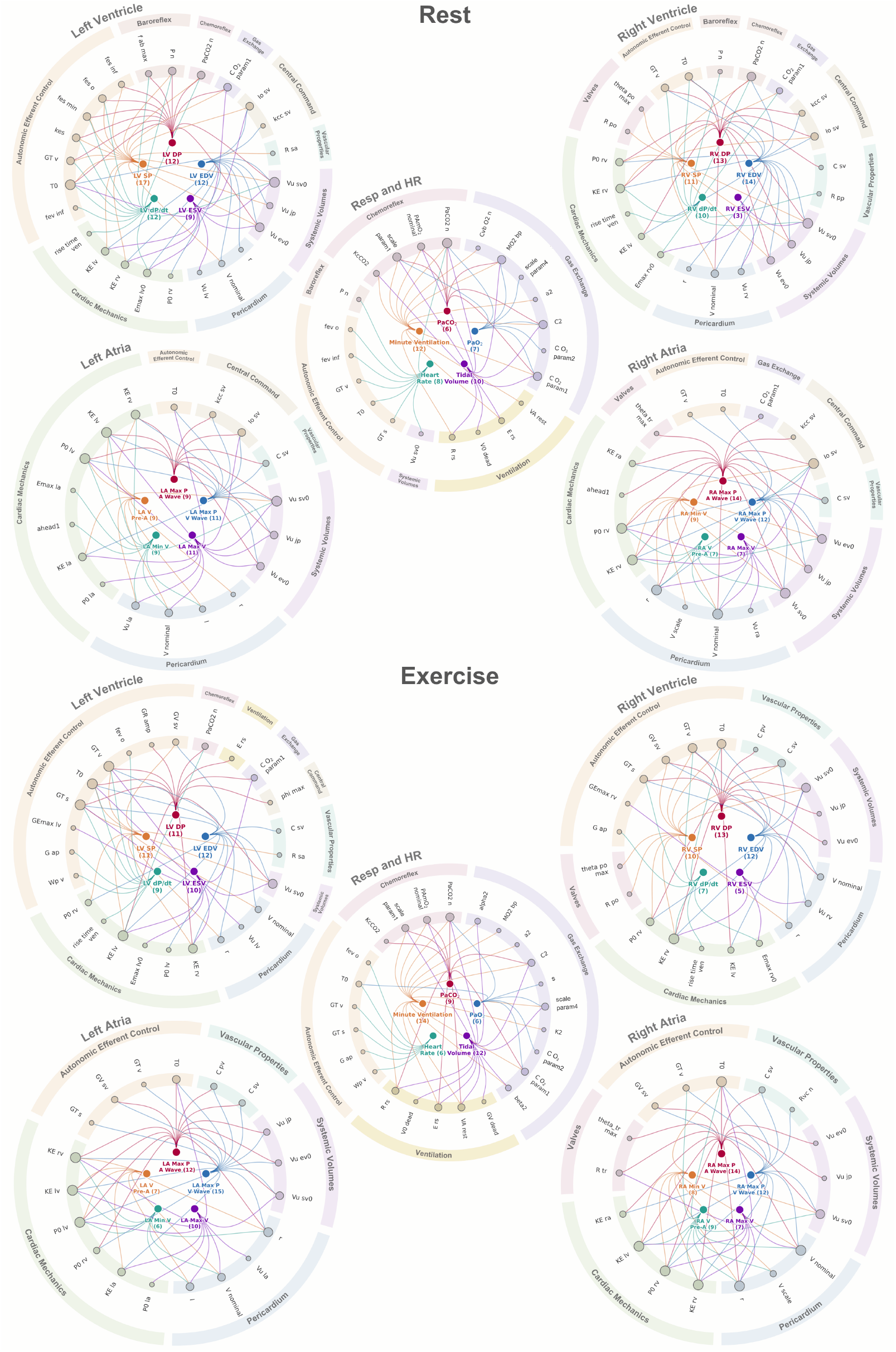
Parameter-target relationships across rest and exercise. Influential parameters are organised by physiological mechanism with groupings for LV, RV, LA, RA, and remaining cardiorespiratory variables. Node size indicates the number of outputs (1 to 5) for which a parameter was identified as influential.

Dominant sensitivities were physiologically coherent. At rest, both ventricular pressure-derivative outputs were influenced by the baseline HR (*T*_0_) and activation timing with ventricular systole (*t*_*rise,ven*_). Minimum ventricular volumes were sensitive to corresponding contractilities (*E*_*max,lv*0*/rv*0_), and unstressed volumes (*V*_*u,lv/rv*_), and maximum ventricular volumes were sensitive to preload-related unstressed volume parameters (*V*_*u,sv*0_). LV pressures were driven by the baroreceptor set-point pressure (*P*_*n*_), whereas RV pressure sensitivities were more distributed across multiple subsystems. Atrial outputs showed clear left-right symmetry, with LA sensitivities clustered around left-sided ventricular passive filling and pericardial parameters and RA outputs dominated by corresponding right-sided quantities (*K*_*E,lv/rv*_, *K*_*E,la/ra*_, *P*_0,*lv/rv*_, *l/r*). All respiratory outputs were sensitive to the nominal PaCO_2_ set point governing chemoreceptor afferent firing (*PaCO*_2,*n*_), and parameters characterising ventilatory mechanics and gas exchange.

With exercise, sensitivities shifted towards autonomic efferent control. Sympathetic and vagal gains modulating heart period, resistances, venous volume, and ventricular contractility (*G*_*T,s*_, *G*_*T,v*_, *G*_*T,sv*_, *G*_*Emax,lv/rv*_) were more influential across ventricular and respiratory targets. Atrial targets became strongly coupled, particularly in the RA, where all five RA targets were sensitive to eight parameters (*K*_*E,lv/rv*_, *P*_0,*rv*_, *R*_*tr*_, *V*_*u,sv*0_, *V*_*nominal*_, *r*, and *T*_0_).

The presence of cardiovascular and control parameters among respiratory sensitivities, and of gas-exchange parameters among cardiovascular sensitivities, reveals the model’s coupled structure.

### 3.2 History matching

After eight waves, 80% training subsets comprising 1054–2612 simulations per wave after simulator-based filtering were sufficient to fit each emulator, achieving a median test *R*^2^ of 0.921 across all 50 targets in the final wave (S4 Appendix, Section 1, Fig A), and a median emulator predictive uncertainty below 10% of the observation uncertainty (S4 Appendix, Section 2). Of the 200,000 parameter sets evaluated per wave, the proportion classified as NROY rose from 4.54% in wave 1, to 27.35% in wave 3, and plateaued near 32% from wave 5 onwards, reflecting the progressive elimination of implausible regions, and successive waves concentrating in the NROY parameter space.

The wave 8 NROY samples defined the prior for subsequent MCMC. Fig 8 portrays the one-dimensional parameter marginals of the NROY sample with the fitted logspline marginals used in the copula prior. In this high-dimensional setting, HM did not narrow individual parameter bounds, but redistributed plausible density within them. Several marginals remain broad, but many are non-uniform, with visible skewness and boundary concentration. Overall, HM focused the emulator training and prior parameter distribution on physiological regions of the parameter space consistent with the data.

**Fig 8.**
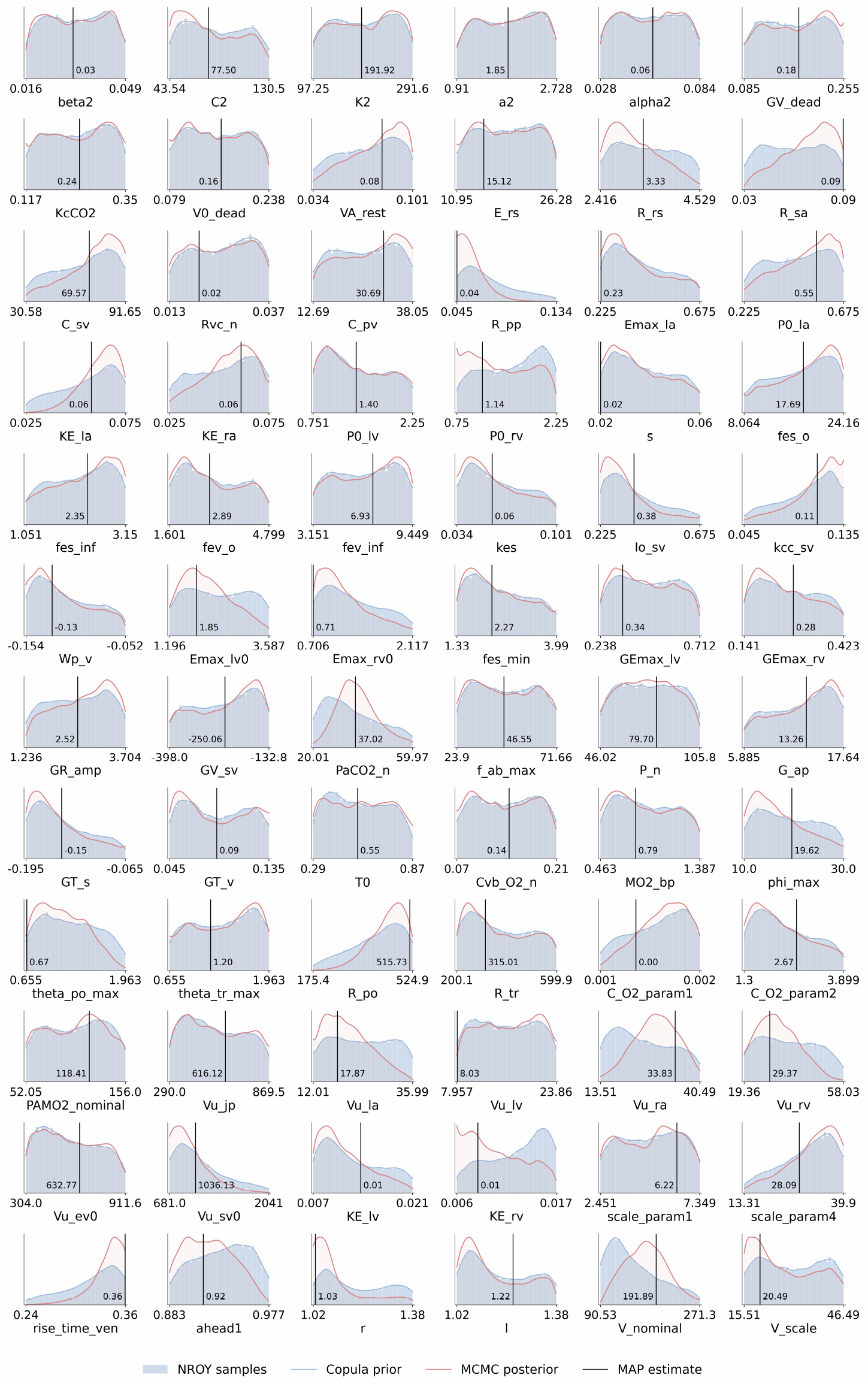
Marginal parameter distributions from History Matching and MCMC. Parameter distributions of the wave 8 NROY parameter sets (blue discrete) overlaid with the fitted logspline marginals defining the copula prior input for MCMC (blue line), and the marginalised posterior distributions after MCMC (red). Vertical lines mark the corresponding parameter value in the maximum a posteriori estimate, which maximises the log posterior and provides the best overall fit to the target data.

### 3.3 MCMC

Four NUTS chains were run with 500 warm-up iterations and 3000 retained post-warmup samples per chain, giving 12,000 posterior draws. Across all calibrated parameters, the maximum split 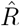 was 1.0010, well below the recommended 1.01 threshold and implying no evidence of between-chain nonconvergence [84]. The minimum effective sample size (ESS), 4738, exceeded the rule-of-thumb of at least 100 effective samples per chain [84]. Thus, for even the slowest-mixing parameter, the 12,000 correlated MCMC draws carried approximately the same information as 4738 independent samples.

The discrete wave 8 NROY parameter sets were fitted with smooth logspline marginals (the prior) and compared with the MCMC marginal posteriors (Fig 8). Posteriors remained broad, signalling weak practical identifiability from the literature targets alone. Some parameters nevertheless display visible prior-to-posterior updating, with posterior contraction in parameters such as *PaCO*_2,*n*_, *V*_*u,la*_, and *E*_*max,lv*0_. The MAP estimate differed from some marginal peaks (e.g. *K*_*E,lv*_, *C*_*O*2,*param*1_), indicating that dense NROY regions do not necessarily correspond to high posterior probability. It also lay on the prior bounds for *R*_*pp*_, *E*_*max,rv*0_, *t*_*rise,ven*_, *s, R*_*sa*_, and *E*_*max,la*_ suggesting that while MCMC improves agreement with the data, the ±50% parameter ranges may truncate the high-posterior region.

Posterior predictive distributions were narrower and closer to the literature targets than both the initial LHC simulations and the wave 8 NROY predictive distributions, with width reductions exceeding 25% for 46 targets (Fig 9). The MAP emulator predictions agreed closely with the corresponding simulator evaluations, with a mean relative difference of 5.1% across the 50 targets and a maximum of 15.0% for the max RA pressure in the V wave (S3 Appendix, Fig G, panel A, Table C). This agreement between the MAP emulator prediction and the corresponding simulator evaluation supports the emulator’s reliability within the calibrated region, consistent with its high performance across targets (median *R*^2^ *>* 0.9).

**Fig 9.**
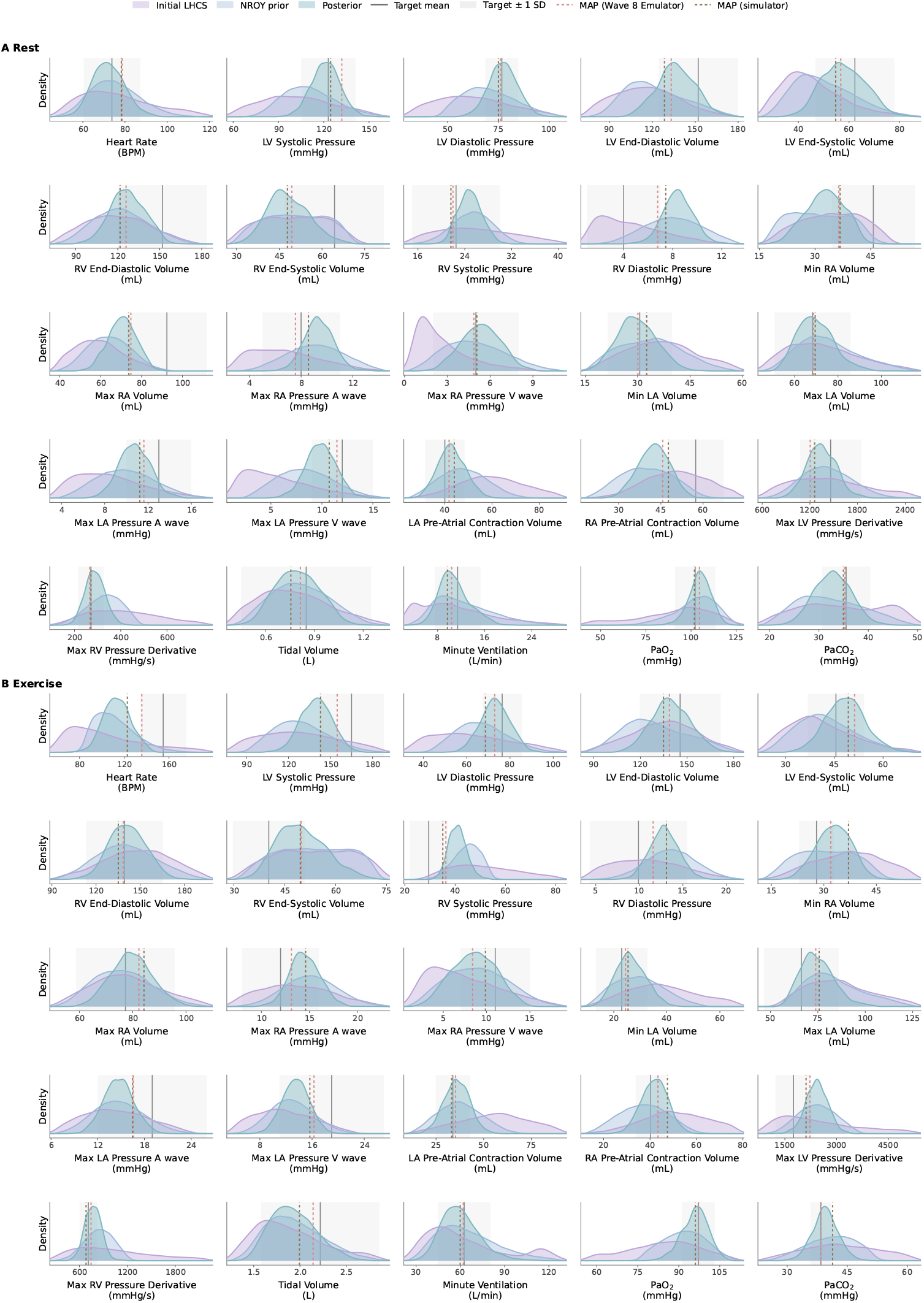
Prior and posterior distributions for calibration targets. Distributions at rest (A) and with exercise (B) are plotted over the central 99.5% range. The three KDEs convey: initial simulator outputs from 8192 LHC parameter sets sampled within ±50% of the nominal parameter set, wave 8 emulator predictions for 63721 NROY parameter sets retained after eight History Matching waves, and the posterior predictive distribution from 1,500 MCMC samples. The black line and shaded band mark the target mean ± 1 observational SD. Dashed lines compare the MAP emulator prediction with the simulator evaluation. 49 of 50 outputs fit within one observational SD, compared to 45 of 50 through the simulator.

Total blood volume (*V*_*tot*_), was fixed rather than inferred as a free parameter because, under mass conservation, *V*_*tot*_ perturbations act as a global stressed-volume offset that confounds with compartmental unstressed volumes. S3 Appendix, Section 5 demonstrates that including *V*_*tot*_ dominated the DGSM ranking without improving agreement with the calibration targets.

### 3.4 Simulator evaluation at the MAP estimate

At rest, the MAP estimate reached a stable periodic state after 690 s for the coupled cardiovascular, respiratory, and gas-exchange system (Fig 10A). The systemic arterial pressure trace displayed the expected systolic upstroke and diastolic runoff, while the ventricular pressure-volume loops preserved left-right physiological differences. The LV loop generated high-pressure systemic ejection, whereas the RV loop operated at a lower pressure while maintaining comparable stroke volume. Ventricular dP/dt was larger in the LV than the RV. Atrial pressure-volume traces formed figure-of-eight loops with clear a- and v-waves and a smaller c-wave-like deflection after atrioventricular valve closure, capturing reservoir, conduit, and booster-pump function. HR, minute ventilation, and arterial gas partial pressures exhibited breath-synchronous oscillations. Overall, the rest-simulated MAP estimate showed strong agreement, with 24 of 25 outputs within 1 target SD and all resting targets within 1.14 target SDs (S3 Appendix, Tables A and B).

**Fig 10.**
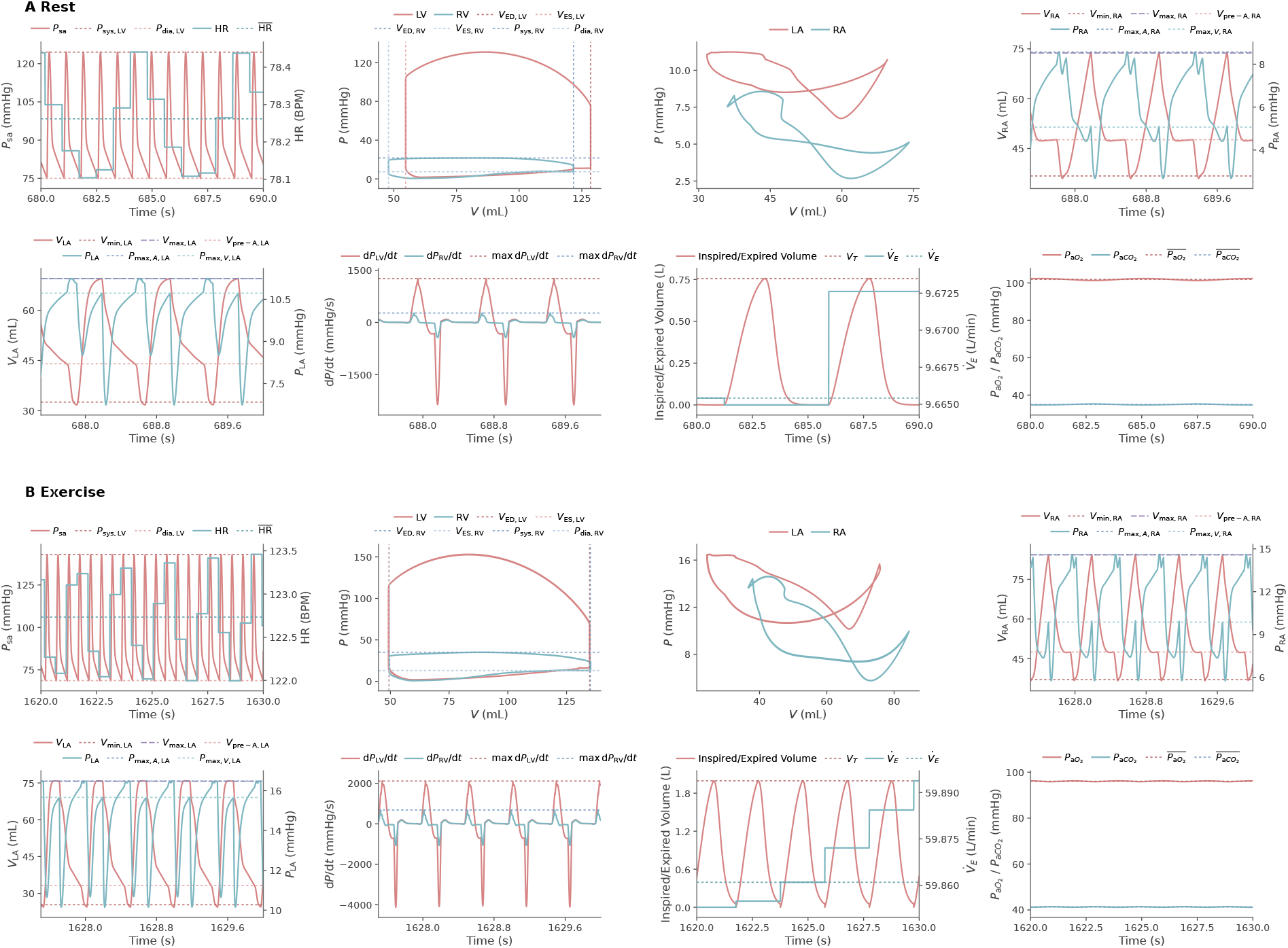
Simulated cardiovascular, respiratory, and gas-exchange dynamics at the MAP estimate. Each panel is a time trace or pressure-volume loop from which the calibration targets were extracted (dashed lines). By 690 s and 1630 s, the system has reached a periodic limit cycle at rest and after exercise stimulation, respectively. Exact output values are reported in S3 Appendix, Section 5, Table C.

With exercise stimulation, metabolic and ventilatory drives are elevated (Fig 10B). Alveolar ventilation rose as the respiratory controller shifted along its work-minimising timing curves (Fig 6B) towards shorter inspiratory and expiratory times. Minute ventilation consequently increased from 9.7 L/min to 59.9 L/min and tidal volume from 0.756 L to 1.99 L, aligning with reported exercise responses [73]. Greater tidal volume increased pulmonary stretch receptor firing, while higher ventilation increased respiratory neuromuscular drive. The modelled decrease in PaO_2_ and increase in PaCO_2_ further enhanced chemoreflex input. Cardiovascular control adapted in parallel. Physiologically, higher arterial pressure raises baroreceptor firing, but central command resets the baroreflex upward and rightward to regulate around a higher operating pressure [55]. In the model, baroreceptor firing oscillated with a greater amplitude during exercise, continuing to reach its upper ceiling each beat but spending more time near its lower bound, slightly reducing time-averaged firing (Fig 11A). The combined afferent inputs amplified cardiac, peripheral, and venous sympathetic outflow (Fig 11B), driving the efferent cardiovascular responses (Fig 11C–F) [55]. Contractility and HR exhibited the expected increases that would elevate stroke volume and cardiac output with exercise [85]. Exercise also redistributes blood volume and vascular resistance. Recruitment of capillary beds shifted unstressed volume into stressed volume, increasing the effective circulating volume. Vasoconstriction in the extrasplanchnic and splanchnic beds increased peripheral resistance and redirected blood flow towards the heart and active muscles, whereas vasodilation reduced resistance and increased perfusion. Overall, these coupled feedback mechanisms rebalanced ventilatory and cardiovascular function to meet the increased metabolic demand.

**Fig 11.**
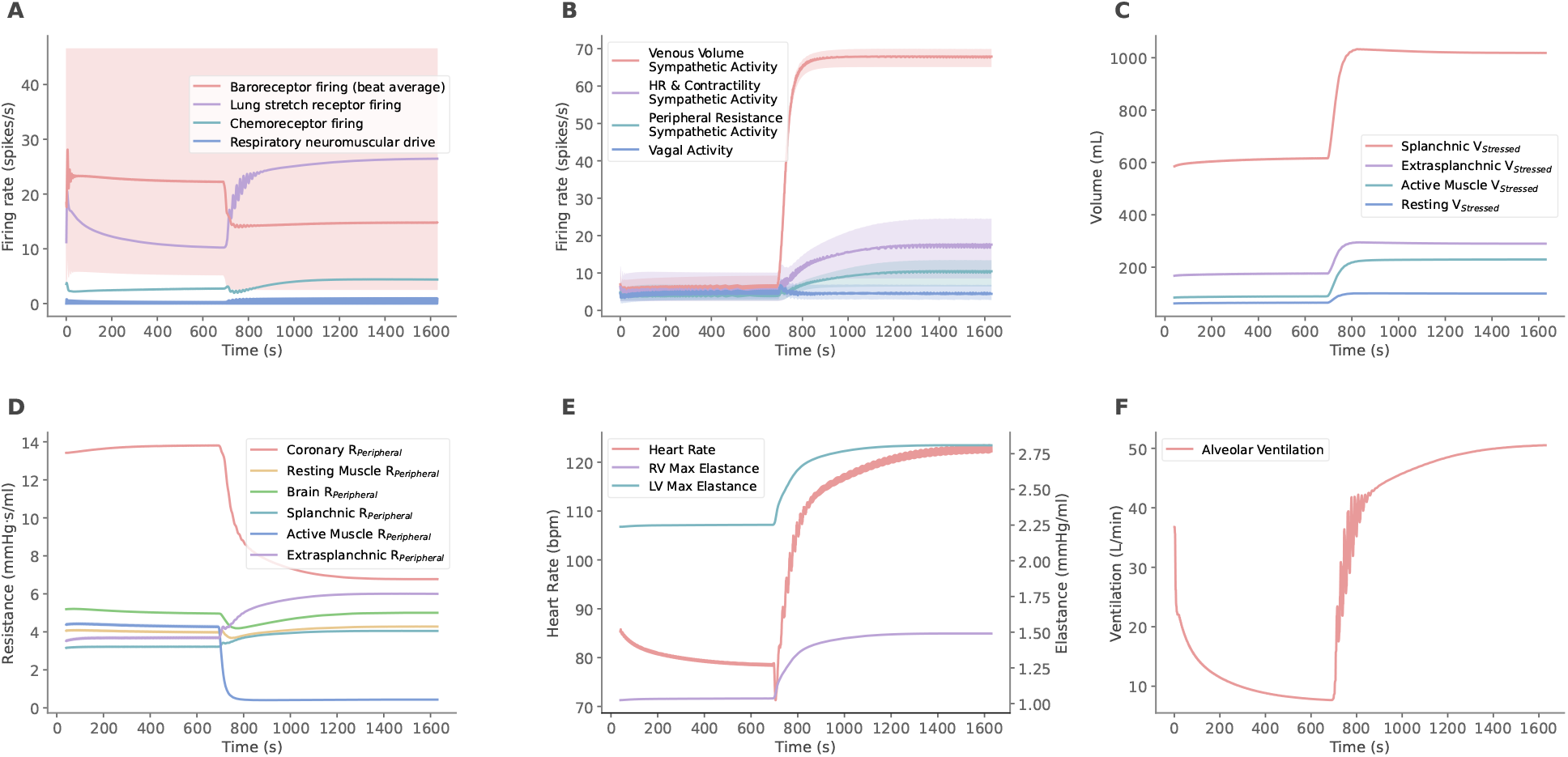
Controlled responses across the rest to exercise transition at 690 s. A: Afferent activity and respiratory neuromuscular drive. B: Autonomic efferent activity. C: Stressed volumes. D: Resistances. E: Heart rate and elastances (contractility). F: Ventilation. Beat-averaged baroreceptor and efferent activity is superimposed on the instantaneous signal (reduced opacity).

### 3.5 Sobol GSA

A constrained emulator-based Sobol GSA was performed in the physiologically plausible 72-parameter space to compare dominant parameters with the DGSM [81]. For a direct comparison, DGSM was recomputed using the simulator for the same 72 parameters at five hundred randomly sampled NROY points. DGSM averages squared local gradients from one-at-a-time simulator perturbations, whereas conditional Sobol total-order effects (*S*_*T*_) quantify output variance under dependence-preserving changes evaluated with the emulator. Thus, DGSM was interpreted as a first-stage screening measure rather than a rank-equivalent estimate of *S*_*T*_ [81]. Expanded *S*_*T*_ and *S*_*i*_ results are reported in S3 Appendix, Section 6, Figs I and J.

Methods broadly agreed across the 50 rest and exercise targets. The mean Spearman rank correlation between the DGSM and *S*_*T*_ profiles over the 72 parameters was 0.79, and 44 of 50 outputs shared at least three of their top five most influential parameters (Fig 12). The union of parameters identified by both methods accounted for a median of 90.5% of the displayed DGSM sensitivity coverage and 96.5% of the displayed *S*_*T*_ coverage. A prominent disagreement occurred for exercise RA V_min_, for which *K*_*E,rv*_ ranked first under DGSM and *V*_*u,ra*_ under conditional *S*_*T*_. Negative, non-physiological volumes with *K*_*E,rv*_ perturbation comprised 10.2% of the retained finite-difference pairs but contributed 48.7% of the summed squared-derivative term, indicating that steep local simulator responses inflated the DGSM ranking (S3 Appendix, Section 6, Fig H). DGSM identified largely the same sensitivity structure as the more computationally demanding Sobol analysis, while remaining differences may reflect the different sensitivity definition or emulator approximation.

**Fig 12.**
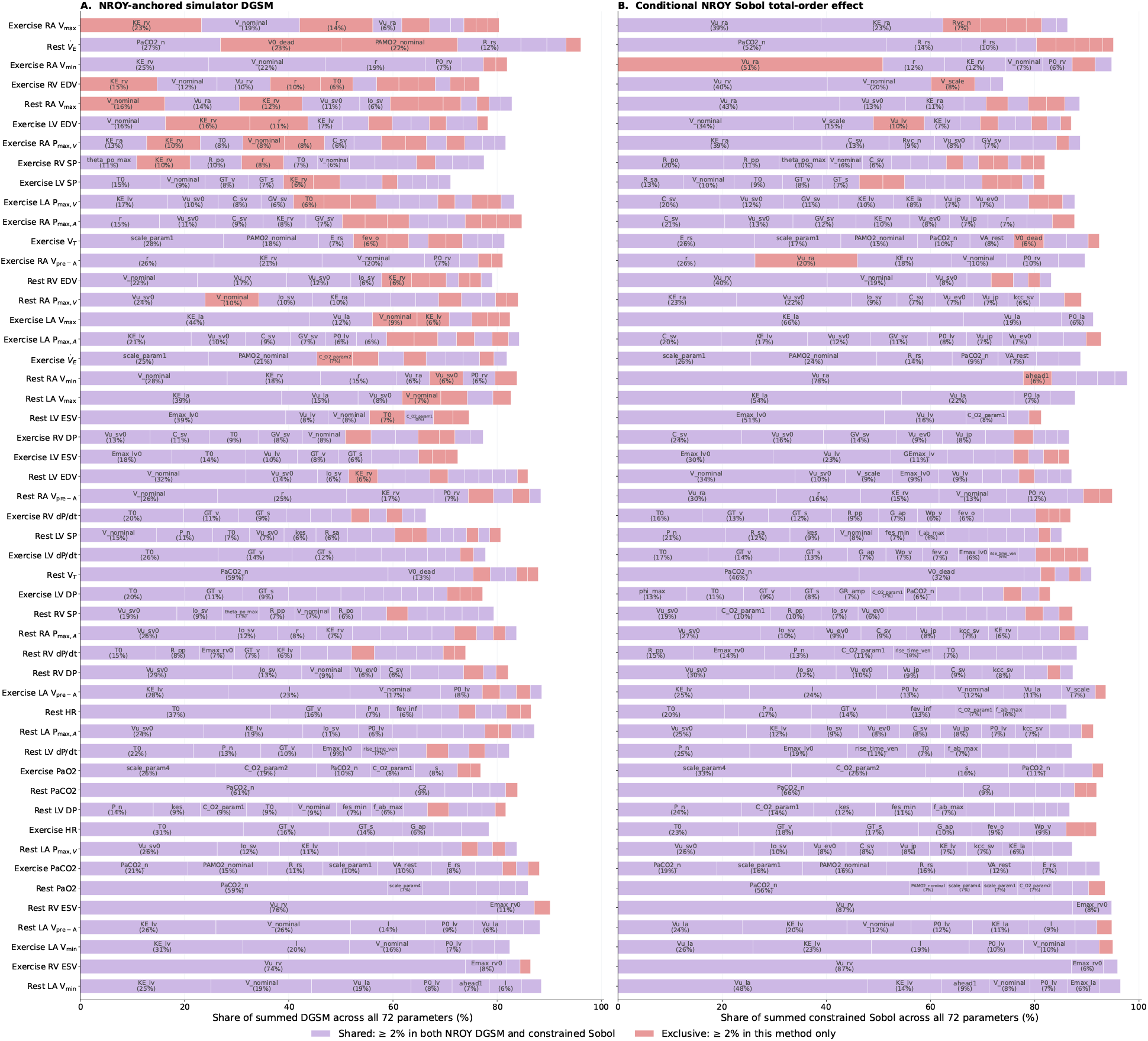
Agreement between simulator-based DGSM and constrained emulator-based Sobol total-order effects. Agreement is shown across 50 rest and exercise targets in the reduced 72 parameter subspace. A: DGSM was calculated using simulator perturbations around base points sampled from the NROY region. B: Conditional *S*_*T*_ was estimated from emulator evaluations over the corresponding NROY distribution. Indices were normalised across all 72 parameters, and the parameters that contribute *>*2% of the summed sensitivity indices for each target are displayed. Targets are ordered from highest to lowest proportion of sensitivity share for the method-exclusive parameters.

## Discussion

The current study demonstrates that a 272-parameter, feedback-regulated cardiorespiratory model can be calibrated simultaneously to 50 rest and exercise literature-derived targets with uncertainty while preserving physiologically coherent dynamics. The work integrates advances in breathing control, atrial dynamics, exercise baroreceptor set point resetting, and physiological feedback delays with a scalable calibration pipeline. The pipeline consists of a simulator-based GSA, HM, MCMC using a Gaussian copula prior fitted to NROY samples, and a validating Sobol GSA constrained to the resulting correlated, non-rectangular domain.

### 4.1 GSA physiological insights and exercise responses

Despite the model’s high dimensionality, only 2–16 parameters influenced each target, with exercise redistributing their relative importance across interacting autonomic, cardiovascular, vascular, and respiratory mechanisms (Fig 12, S3 Appendix, Section 6, Fig I). At rest, volume targets were sensitive to both chamber-specific unstressed volumes and cardiac mechanics. Unstressed volumes dominated summed conditional *S*_*T*_, whereas LV ESV and LA V_max_ were most sensitive to contractility *E*_*max,lv*0_ (51.1%) and passive stiffness *K*_*E,la*_ (54.5%), respectively. For LV SP, the baroreceptor set point *P*_*n*_ contributed 20.6% of the summed conditional *S*_*T*_, while HR was most influenced by the baseline heart period *T*_0_ (20.4%), *P*_*n*_ (16.7%), vagal heart period gain *G*_*T,v*_ (14.4%), and maximum vagal efferent firing, *f*_*ev,inf*_ (13.4%). This agrees with Tlalka et al. (2024), whose regulated resting models were most sensitive to *P*_*n*_, *G*_*T,v*_, and *f*_*ev,inf*_ [4]. During exercise, influence shifted further towards regulated parameters, with *T*_0_, *G*_*T,v*_ and sympathetic heart period gain, *G*_*T,s*_, accounting for larger shares of HR, ventricular pressures, and pressure derivatives than at rest. These rankings emerged dynamically after exercise onset. Sympathetic outflow increased, coordinating increases in HR, contractility, stressed volume, and resistance in non-exercising vascular beds (Fig 11). This is consistent with experimental evidence of arterial baroreflex resetting to sustain pressures as vagal modulation gives way to sympathetic dominance, and with models where active-muscle vasodilation, splanchnic-muscle vasoconstriction, and central command reproduce exercise responses [17, 19, 23, 86–88]. Respiratory targets showed similar homeostatic feedback through the chemoreflex, particularly by basal PaCO_2_ (*PaCO*_2,*n*_). A rise in PaCO_2_ or fall in PaO_2_ increased chemoreceptor firing and alveolar ventilation. Greater airflow across the lungs increases O_2_ uptake and CO_2_ removal, opposing the disturbance and keeping gas partial pressures near resting values despite the substantial rise in metabolic demand during exercise [17, 89]. Exercise therefore redistributed influence across a coupled network of autonomic and respiratory control pathways. Rapid autonomic and haemodynamic adjustments at exercise onset overlap the slower ventilatory, CO_2_-storage, and chemoreflex processes, producing responses spanning seconds to several minutes [90, 91].

Beyond calibration targets, the model reproduced physiological integrated exercise responses that were not explicitly calibrated for. As the imposed oxygen consumption 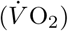 increased from 0.25 to 1.20 L/min, cardiac output 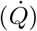 rose from 5.78 to 10.5 L/min, giving a 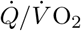 of 4.97 L/L, which is within the broader reported range of 4.6 to *>*6.0 L/L and slightly lower than the 5.4–5.9 L/L observed in male endurance-trained athletes during submaximal cycling [92]. This reflected a 57% increase in HR and ventricular volume shifts that increased stroke volume by 16% and ejection fraction from 57.4% to 63.4%. Although modest, these changes were physiological and consistent with healthy responses at comparable 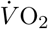 (17.5 mL kg^−1^ · min^−1^ = 1.2 L · min^−1^ at 70 kg) [85, 93]. By the Fick principle, the arterial-venous oxygencontent difference 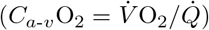 widened from 4.21 to 11.5 mL/dL, indicating greater peripheral oxygen extraction and agreeing with the exercise-induced fall in mixed-venous oxygen saturation [94]. Moreover, as tidal volume increased from 0.756 to 1.99 L, dead-space volume increased from 0.184 to 0.312 L, aligning with graded exercise studies [95]. The separate alveolar and arterial compartments and explicit pulmonary shunt also enabled estimation of the alveolar-arterial oxygen widening from 4.2 to 12.3 mmHg as exercise arterial pO_2_ decreased, consistent with the exercise-induced widening reported in healthy adults [96]. Further calibration across multiple exercise workloads could enable comparison with CPET-derived trajectories, slopes, and thresholds [97]. Together, these results show that the coupled cardiovascular and gas-exchange formulations reproduce expected rest-to-exercise responses beyond the quantities used for calibration.

### 4.2 DGSM and Sobol GSA

The preliminary GSA functions to isolate mechanisms in a highly coupled and dimensional model so subsequent fitting is restricted to parameters that materially influence the observed data. Simulator-based DGSM GSA converged in 250 base points, corresponding to 250(272 + 1) = 68,250 simulator evaluations (S3 Appendix, Section 3). In contrast, published Sobol GSAs required 7.95 million direct model evaluations for a regulated 51-parameter four-chamber model and 94,000 GPE evaluations for a reduced 45-parameter whole-heart model, highlighting the prohibitive cost of direct Sobol analysis on a 272-parameter simulator [4, 34]. An emulator-based GSA would additionally require accurate surrogates across a high-dimensional space, a challenge complicated by delays, closed-loop feedback, sharp transitions, and instability boundaries. Simulator-evaluated DGSM retains these behaviours and exposes numerical fragility that emulator smoothing obscures (S3 Appendix, Section 6), while efficiently providing converged influential parameters for subsequent calibration. This practically balances biophysical fidelity and computational cost for population-level studies [98].

Variance-based indices are defined relative to the assumed input distribution, so their values and rankings depend on the domain over which they are computed [99, 100]. We therefore constrained Sobol GSA over the final NROY region rather than the full pre-HM prior, 96% of which was physiologically implausible (Section 3.2). Working within this correlated NROY domain also changes how the indices are interpreted. *S*_*T*_ may be smaller than *S*_*i*_, the indices need not sum to one, and their difference does not represent interaction [81]. For rest EDV, *V*_*nominal*_ *S*_*i*_ = 0.695 but conditional *S*_*T*_ = 0.153 (S3 Appendix, Section 6, Figs I and J). Its correlations with other EDV-sensitive parameters (*K*_*E,lv*_, *r, R*_*po*_, and *K*_*E,rv*_; |*r*| = 0.28 to 0.41) are captured by *S*_*i*_ but largely removed from *S*_*T*_ once those parameters are fixed (S3 Appendix, Section 6, Fig K). The normalised indices rank relative influence within the fitted wave 8 NROY distribution rather than partitioning output variance as in classical independent-input Sobol analysis. These dependencies may also contribute to the *S*_*T*_ and DGSM difference. DGSM averages one-at-a-time local gradients while in estimating *S*_*T*_, *X*_*i*_ variability is restricted once *X*_−*i*_ is fixed [81]. Parameter correlations can significantly affect SA results, highlighting the need to assess input correlations during model development [82].

### 4.3 Mechanistic limitations

We aimed to develop a more comprehensive and physiologically grounded model of cardiorespiratory function and regulation at rest and during exercise, while retaining computational efficiency and tractability through necessary simplifying assumptions. Our time-varying elastance model represents active and passive chamber pressures but not fibre-level myocardial stresses or explicit interventricular septal mechanics [6,101]. Currently, interventricular interaction is characterised indirectly through closed-loop circulation, where RV output determines subsequent LV filling, and a shared pericardial pressure derived from total four-chamber volume. Accordingly, passive diastolic stiffnesses *K*_*E,rv*_ and *K*_*E,lv*_, and pericardial parameters, together influenced multiple pressure and volume targets (Fig 7). Among the larger model-observation residuals were exercise RA V_min_ and RA V_pre-A_, which were 1.83 and 1.12 SD above their respective targets and were sensitive to *K*_*E,rv*_ and pericardial parameters, suggesting incomplete representation of right-sided filling mechanics. During exercise, diastolic ventricular interaction can also become clinically significant, so septal effects may no longer be negligible [102]. A Laplace-based fibre model would be valuable for studying ventricular loading, remodelling, or dyssynchrony [34, 103].

Other model simplifications improve scalability for large-scale calibration and sensitivity analysis, but limit applications where respiratory mechanics, posture, or cellular physiology are central. The model neglects the effects of the muscle pump and intrathoracic and abdominal pressure oscillations on cardiovascular function. Because targets were averaged over the final 10 heart beats, finite-difference estimates could be sensitive to respiratory oscillations and spuriously inflate DGSM values. Sarmiento et al. (2021) reported only small haemodynamic changes after excluding the respiratory and muscle pumps, but Angus et al. (2023) found that attenuating intrathoracic-pressure swings reduced exercise cardiac output [17, 104]. Restoring these mechanisms could improve our underpredicted exercise HR and also account for physiological respiratory sinus arrhythmia effects [105]. Omitting gravity likewise prevents characterisation of lower-body pooling and reduced preload, so hydrostatic venous distension and lower-body baroreceptor responses should be modelled for upright simulations [106]. The present phenomenological baroreflex also cannot capture the ionic current dynamics at the afferent baroreflex limb and NTS neuron represented by Hodgkin-Huxley-type models [27].

### 4.4 Parameter calibration

MCMC reweighted the wave-8 NROY region for simultaneous agreement with 50 targets. The central 95% of posterior samples spanned at least 90% of the original parameter range for 48 of 72 parameters (Fig 8), indicating weak practical identifiability. Pairwise dependence was sparse overall, but several relationships strengthened relative to the HM distribution, including *V*_*nominal*_–*V*_*scale*_, *K*_*E,lv*_–*l*, and *E*_*max,lv*0_–*V*_*u,lv*_ (Fig 13). These dependencies suggest parameter compensation, where multiple combinations reproduce the targets, explaining the limited contraction of individual marginals [107]. Yet, all target distributions narrowed relative to the NROY prior (Fig 9). Broad parameter posteriors can still support task-specific predictions, and narrow output intervals do not require every parameter to be precisely estimated [108, 109]. The targets therefore constrain parameter combinations rather than individual values, from which the MAP estimate provided the single best-fitting parameter set for both rest and exercise.

**Fig 13.**
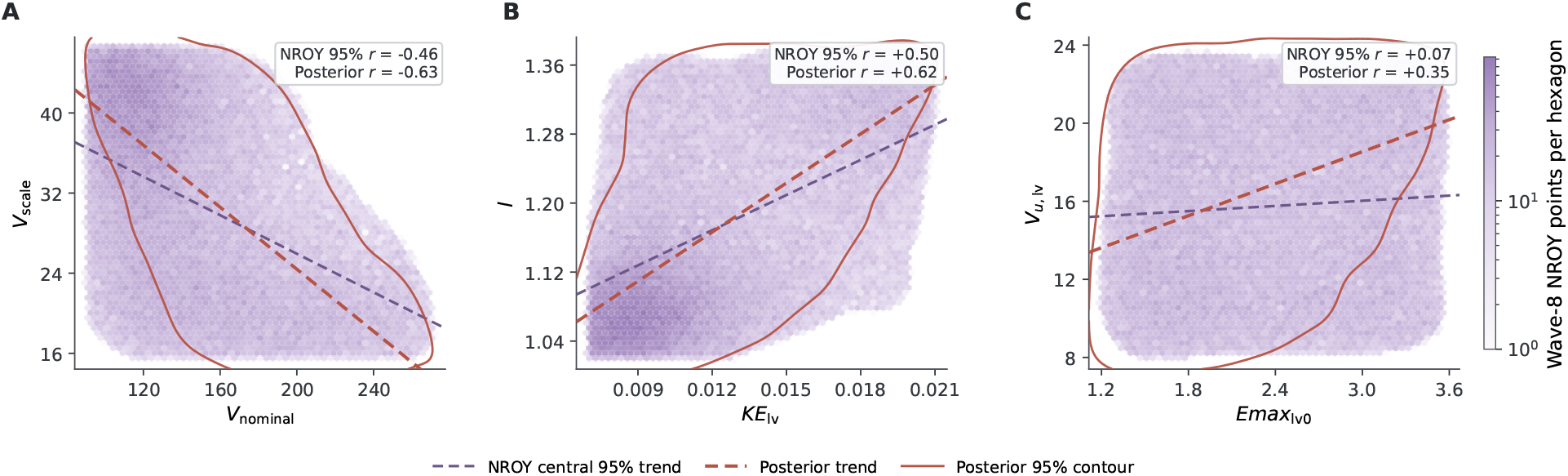
Selected pairwise parameter dependencies within the wave-8 NROY region and MCMC posterior. A: *V*_*nominal*_–*V*_*scale*_. B: *K*_*E,lv*_–*l*. C: *E*_*max,lv*0_–*V*_*u,lv*_. Hexagons show the central 95% wave-8 NROY point density, contours enclose the smoothed 95% highest-density regions of the 12,000 posterior samples, and dashed lines show the corresponding least-squares trends.

Emulator-based MCMC made calibration computationally feasible, and the MAP predictions agreed closely between the emulator and the full simulator (Fig 9). The emulator-simulator difference implies emulator error so the posterior may differ modestly from the full-simulator posterior [110]. Target distributions (Fig 9) used only emulator mean predictions and only represent output uncertainty from the plausible posterior parameter sets. Although GPE predictive variance was combined with target variance in the likelihood (Eqs (21) and (22)), no model-discrepancy term was included (Eq (19)). The posterior was therefore conditional on the assumed model structure and emulator accuracy. An extension would be to model simulated-target mismatch explicitly using GPs, or use multi-output GPEs where target residuals are correlated [109, 111]. Emulator reliability could also be improved by adding simulator runs in plausible parameter regions where they are expected to reduce uncertainty in the estimated posterior [112]. This would help distinguish between residuals from emulator error vs omitted physiology.

The same MAP parameter set reproduced both states, so rest and exercise residuals could not be minimised independently and instead exposed the model’s structural capacity to span both. Exercise HR was 1.54 SD below its target whereas maximum LV dP/dt was 0.70 SD above. These opposing residuals may indicate autonomic regulation of heart period and ventricular elastance without explicit HR-dependent modulation of peak elastance via the force-frequency response [113]. However, because these targets were population summaries, residuals may instead point to incompatible joint target values rather than model limitations.

The present joint calibration of rest and exercise extends previous Bayesian calibration of LPMs, which typically fitted one physiological state at a time and estimated smaller parameter sets against fewer summary targets [38, 43, 109, 114]. These studies combined initial optimisation with adaptive or gradient-based MCMC. Fit quality depended strongly on target construction, with greater uncertainty assigned to low-confidence targets, allowing larger residuals [38, 43, 109]. Exercise studies, moreover, have relied on manually tuned or error-minimised point estimates fitted to observed or population data [29, 30, 115, 116]. Point estimates quantify neither joint parameter nor predictive uncertainty, concealing alternative parameter combinations and mechanisms consistent with the data. By inferring a joint posterior across both states from population summary targets, the present study provides an uncertainty-quantified population prior. Sequential updating with longitudinal patient data, so that each posterior informs the prior of the next, would refine the population representation to reflect the individual, the foundation for a cardiovascular digital twin [37, 38].

## Conclusion

We developed OpenCRS, a 272-parameter, feedback-regulated cardiorespiratory model with a scalable workflow for global sensitivity analysis and joint rest-exercise calibration. Simulator-based DGSM reduced 272 parameters to 72 across 50 targets and recovered the dominant sensitivities identified by constrained Sobol analysis within the correlated, physiologically non implausible region. History Matching and MCMC identified parameter combinations consistent with the literature-derived targets and narrowed predictive distributions. The MAP parameter set represented both states through the model’s embedded feedback mechanisms rather than through independent rest and exercise fitting. The posterior therefore provides an informed population-level prior, while the model offers a modular basis for stimuli-specific sensitivity analysis. Its value lies not only in prediction, but in linking physiological changes to underlying mechanisms.

## Data Availability

No new experimental or participant data were generated. Calibration targets were obtained from published sources cited in the manuscript. The OpenCRS model and code for performing the global sensitivity analyses, training emulators, conducting History Matching and MCMC, and generating the figures, are publicly available at https://github.com/Sheng-Ya/OpenCRS under the MIT Licence. The numerical data underlying the results are archived on Zenodo at https://doi.org/10.5281/zenodo.22732021 under a CC BY 4.0 licence.

## Supporting information

### S1 Appendix. Model equations and parameters

Complete equations and nominal parameter values for the cardiovascular system and controller, respiratory controller and mechanics, breathing-pattern optimiser, gas exchange and transport, and metabolic dynamics, together with justifications for altered parameter values. Contains Figs A and B and Tables A–AB.

### 2 Appendix. State variable initial conditions

Initial values for the 82 OpenCRS state variables: 25 cardiovascular, 32 cardiovascular-control, 24 gas-exchange, and one respiratory-control state, together with the gas-store initialisation procedure. Contains Table A.

### S3 Appendix. Derivative-based global sensitivity measure

SALib DGSM function edits (Section 1), robustness to parameter bounds (Section 2), base-point convergence (Section 3), expanded DGSM results for individual targets (Section 4), total blood volume sensitivity and calibration analyses (Section 5), and comparison of DGSM with constrained Sobol analysis (Section 6). Contains Figs A–K and Tables A–C.

### S4 Appendix. History Matching and MCMC

Emulator accuracy across History Matching waves, including test *R*^2^ by target and the local limitation for rest minute ventilation (Section 1), and emulator predictive variance relative to observation variance (Section 2). Contains Figs A–C.

## Author Contributions

**Conceptualization:** Sheng-Ya Wang, Harry Saxton, Maximilian Balmus, Steven A. Niederer

**Data Curation:** Sheng-Ya Wang

**Formal Analysis:** Sheng-Ya Wang

**Funding Acquisition:** Steven A. Niederer

**Investigation:** Sheng-Ya Wang

**Methodology:** Sheng-Ya Wang, Harry Saxton, Maximilian Balmus, Steven A. Niederer

**Project Administration:** Sheng-Ya Wang, Steven A. Niederer

**Resources:** Steven A. Niederer

**Software:** Sheng-Ya Wang

**Supervision:** Harry Saxton, Maximilian Balmus, Steven A. Niederer

**Validation:** Sheng-Ya Wang

**Visualization:** Sheng-Ya Wang

**Writing–Original Draft Preparation:** Sheng-Ya Wang

**Writing–Review & Editing:** Sheng-Ya Wang, Harry Saxton, Maximilian Balmus, Steven A. Niederer

## References

1. Lwin M, Masding A, McCabe C. Physical exercise for pulmonary arterial hypertension diagnosis and therapy. Int J Cardiol Congenit Heart Dis. 2025 Mar;19:100565. doi:10.1016/j.ijcchd.2025.100565. PMID: 40066343.

2. Houstis NE, Eisman AS, Pappagianopoulos PP, Wooster L, Bailey CS, Wagner PD, et al. Exercise Intolerance in Heart Failure With Preserved Ejection Fraction. Circulation. 2018 Jan;137(2):148–61. doi:10.1161/CIRCULATIONAHA.117.029058. PMID: 28993402.

3. Pironet A, Docherty PD, Dauby PC, Chase JG, Desaive T. Practical identifiability analysis of a minimal cardiovascular system model. Comput Methods Programs Biomed. 2019 Apr;171:53–65. doi:10.1016/j.cmpb.2017.01.005. PMID: 28153466.

4. Tlalka K, Saxton H, Halliday I, Xu X, Narracott A, Taylor D, et al. Sensitivity analysis of closed-loop one-chamber and four-chamber models with baroreflex. PLOS Comput Biol. 2024 Dec;20(12):e1012377. doi:10.1371/journal.pcbi.1012377. PMID: 39715272.

5. Niederer SA, Lumens J, Trayanova NA. Computational models in cardiology. Nat Rev Cardiol. 2019 Feb;16(2):100–11. doi:10.1038/s41569-018-0104-y. PMID: 30361497.

6. Haghebaert M, Varsos P, Meiburg R, Vignon-Clementel I. A comparative study of lumped heart models for personalized medicine through sensitivity and identifiability analysis. J Physiol. 2025 May. doi:10.1113/JP287929. PMID: 40349323.

7. Colli Franzone P, Pavarino LF, Scacchi S. Mathematical Cardiac Electrophysiology. vol. 13 of MS&A. Cham: Springer International Publishing; 2014. doi:10.1007/978-3-319-04801-7.

8. van de Vosse FN, Stergiopulos N. Pulse Wave Propagation in the Arterial Tree. Annu Rev Fluid Mech. 2011;43:467–99. doi:10.1146/annurev-fluid-122109-160730.

9. Walker M, Moore H, Ataya A, Pham A, Corris PA, Laubenbacher R, et al. A perfectly imperfect engine: Utilizing the digital twin paradigm in pulmonary hypertension. Pulm Circ. 2024;14(2):e12392. doi:10.1002/pul2.12392. PMID: 38933181.

10. Saxton H. Uncertainty quantification and personalisation of lumped parameter models of the cardiovascular system [doctoral]. Sheffield Hallam University; 2025. doi:10.7190/shu-thesis-00673.

11. Odeigah OO, Valdez-Jasso D, Wall ST, Sundnes J. Computational models of ventricular mechanics and adaptation in response to right-ventricular pressure overload. Front Physiol. 2022 Aug;13. doi:10.3389/fphys.2022.948936. PMID: 36091369.

12. Gall AL, Vallée F, Pushparajah K, Hussain T, Mebazaa A, Chapelle D, et al. Monitoring of cardiovascular physiology augmented by a patient-specific biomechanical model during general anesthesia. A proof of concept study. PLOS One. 2020 May;15(5):e0232830. doi:10.1371/journal.pone.0232830. PMID: 32407353.

13. Hu Z, Herrmann JE, Schwarz EL, Gerosa FM, Emuna N, Humphrey JD, et al. Multiphysics Simulations of a Bioprinted Pulsatile Fontan Conduit. J Biomech Eng. 2025 May;147(7):071001. doi:10.1115/1.4068319. PMID: 40172060.

14. Richter J, Nitzler J, Pegolotti L, Menon K, Biehler J, Wall WA, et al. Bayesian Windkessel calibration using optimized zero-dimensional surrogate models. Philos Trans A Math Phys Eng Sci. 2025 Mar;383(2292):20240223. doi:10.1098/rsta.2024.0223. PMID: 40078147.

15. Cheng L, Ivanova O, Fan HH, Khoo MCK. An integrative model of respiratory and cardiovascular control in sleep-disordered breathing. Respir Physiol Neurobiol. 2010 Nov;174(1-2):4–28. doi:10.1016/j.resp.2010.06.001. PMID: 20542148.

16. Ishbulatov YM, Karavaev AS, Kiselev AR, Simonyan MA, Prokhorov MD, Ponomarenko VI, et al. Mathematical modeling of the cardiovascular autonomic control in healthy subjects during a passive head-up tilt test. Sci Rep. 2020 Oct;10(1):16525. doi:10.1038/s41598-020-71532-7. PMID: 33020530.

17. Sarmiento CA, Hernández AM, Serna LY, Mañanas MA. An integrated mathematical model of the cardiovascular and respiratory response to exercise: model-building and comparison with reported models. Am J Physiol Heart Circ Physiol. 2021 Apr;320(4):H1235–60. doi:10.1152/ajpheart.00074.2020. PMID: 33416450.

18. Zambrano BA, Gharahi H, Lim C, Jaberi FA, Choi J, Lee W, et al. Association of intraluminal thrombus, hemodynamic forces, and abdominal aortic aneurysm expansion using longitudinal CT images. Ann Biomed Eng. 2016 May;44(5):1502–14. doi:10.1007/s10439-015-1461-x. PMID: 26429788.

19. Magosso E, Ursino M. Cardiovascular response to dynamic aerobic exercise: a mathematical model. Med Biol Eng Comput. 2002 Nov;40(6):660–74. doi:10.1007/BF02345305. PMID: 12507317.

20. Ursino M, Magosso E. Acute cardiovascular response to isocapnic hypoxia. I. A mathematical model. Am J Physiol Heart Circ Physiol. 2000 Jul;279(1):H149–65. doi:10.1152/ajpheart.2000.279.1.H149. PMID: 10899052.

21. Ursino M. Interaction between carotid baroregulation and the pulsating heart: a mathematical model. Am J Physiol Heart Circ Physiol. 1998 Nov;275(5):H1733–47. doi:10.1152/ajpheart.1998.275.5.H1733. PMID: 9815081.

22. Suga H, Sagawa K. Instantaneous pressure-volume relationships and their ratio in the excised, supported canine left ventricle. Circ Res. 1974 Jul;35(1):117–26. doi:10.1161/01.res.35.1.117. PMID: 4841253.

23. Elstad M, Toska K, Walløe L. Model simulations of cardiovascular changes at the onset of moderate exercise in humans. J Physiol. 2002 Sep;543(Pt 2):719–28. doi:10.1113/jphysiol.2002.019422. PMID: 12205203.

24. D’Angelo C, Papelier Y. Mathematical modelling of the cardiovascular system and skeletal muscle interaction during exercise. ESAIM Proc. 2005 Sep;14:72–88. doi:10.1051/proc:2005007.

25. Lumens J, Delhaas T, Kirn B, Arts T. Three-Wall Segment (TriSeg) Model Describing Mechanics and Hemodynamics of Ventricular Interaction. Ann Biomed Eng. 2009 Nov;37(11):2234–55. doi:10.1007/s10439-009-9774-2. PMID: 19718527.

26. Korakianitis T, Shi Y. Numerical simulation of cardiovascular dynamics with healthy and diseased heart valves. J Biomech. 2006;39(11):1964–82. doi:10.1016/j.jbiomech.2005.06.016. PMID: 16140309.

27. Fernandes LG, Müller LO, Feijoó RA, Blanco PJ. Closed-loop baroreflex model with biophysically detailed afferent pathway. Int J Numer Method Biomed Eng. 2024 Sep;40(9):e3849. doi:10.1002/cnm.3849. PMID: 39054666.

28. Sheel AW, Romer LM. Ventilation and Respiratory Mechanics. Compr Physiol. 2012;2(2):1093–142. doi:10.1002/j.2040-4603.2012.tb00417.x. PMID: 23798297.

29. Albanese A, Cheng L, Ursino M, Chbat NW. An integrated mathematical model of the human cardiopulmonary system: model development. Am J Physiol Heart Circ Physiol. 2016 Apr;310(7):H899–921. doi:10.1152/ajpheart.00230.2014. PMID: 26683899.

30. Serna LY, Mañanas MA, Hernández AM, Rabinovich RA. An Improved Dynamic Model for the Respiratory Response to Exercise. Front Physiol. 2018 Feb;9. doi:10.3389/fphys.2018.00069. PMID: 29467674.

31. Păun LM, Qureshi MU, Colebank M, Hill NA, Olufsen MS, Haider MA, et al. MCMC methods for inference in a mathematical model of pulmonary circulation. Stat Neerl. 2018;72(3):306–38. doi:10.1111/stan.12132.

32. Sobol IM, Kucherenko S. A new derivative based importance criterion for groups of variables and its link with the global sensitivity indices. Comput Phys Commun. 2010 Jul;181(7):1212–7. doi:10.1016/j.cpc.2010.03.006.

33. Saltelli A, Annoni P, Azzini I, Campolongo F, Ratto M, Tarantola S. Variance based sensitivity analysis of model output. Design and estimator for the total sensitivity index. Comput Phys Commun. 2010 Feb;181(2):259–70. doi:10.1016/j.cpc.2009.09.018.

34. Strocchi M, Longobardi S, Augustin CM, Gsell MAF, Petras A, Rinaldi CA, et al. Cell to whole organ global sensitivity analysis on a four-chamber heart electromechanics model using Gaussian processes emulators. PLOS Comput Biol. 2023 Jun;19(6):e1011257. doi:10.1371/journal.pcbi.1011257. PMID: 37363928.

35. Peirlinck M, Sahli Costabal F, Sack KL, Choy JS, Kassab GS, Guccione JM, et al. Using machine learning to characterize heart failure across the scales. Biomech Model Mechanobiol. 2019 Dec;18(6):1987–2001. doi:10.1007/s10237-019-01190-w. PMID: 31240511.

36. Martinez ES, Moscoloni B, Salvador M, Kong F, Peirlinck M, Marsden AL. Full-field surrogate modeling of cardiac electrophysiology encoding geometric variability. Comput Methods Appl Mech Eng. 2026 Jan;448:118444. doi:10.1016/j.cma.2025.118444. PMID: 41928767.

37. Corral-Acero J, Margara F, Marciniak M, Rodero C, Loncaric F, Feng Y, et al. The ‘Digital Twin’ to enable the vision of precision cardiology. Eur Heart J. 2020 Dec;41(48):4556–64. doi:10.1093/eurheartj/ehaa159. PMID: 32128588.

38. Thiel JN, Zlicar M, Steinseifer U, Kirn B, Neidlin M. Bayesian parameter inference and uncertainty-informed sensitivity analysis in a 0D cardiovascular model for intraoperative hypotension. Comput Biol Med. 2026 Jan;200:111371. doi:10.1016/j.compbiomed.2025.111371. PMID: 41353885.

39. Korakianitis T, Shi Y. A concentrated parameter model for the human cardiovascular system including heart valve dynamics and atrioventricular interaction. Med Eng Phys. 2006 Sep;28(7):613–28. doi:10.1016/j.medengphy.2005.10.004. PMID: 16293439.

40. Chiari L, Avanzolini G, Ursino M. A comprehensive simulator of the human respiratory system: validation with experimental and simulated data. Ann Biomed Eng. 1997;25(6):985–99. doi:10.1007/BF02684134.

41. Serna Higuita LY, Mañanas MA, Mauricio Hernández A, Marína Sánchez J, Benito S. Novel Modeling of Work of Breathing for Its Optimization During Increased Respiratory Efforts. IEEE Syst J. 2016 Sep;10(3):1003–13. doi:10.1109/JSYST.2014.2323114.

42. Castro R, Kattan E, Retamal J, Hernández G, Pinsky MR. Venous congestion from a vascular waterfall perspective: reframing congestion as a dynamic Starling resistor phenomenon. Intensive Care Med Exp. 2025 Nov;13(1):119. doi:10.1186/s40635-025-00828-7. PMID: 41284176.

43. Schiavazzi DE, Baretta A, Pennati G, Hsia TY, Marsden AL. Patient-specific parameter estimation in single-ventricle lumped circulation models under uncertainty. Int J Numer Method Biomed Eng. 2017 Mar;33(3). doi:10.1002/cnm.2799. PMID: 27155892.

44. Pant S, Corsini C, Baker C, Hsia TY, Pennati G, Vignon-Clementel IE. A Lumped Parameter Model to Study Atrioventricular Valve Regurgitation in Stage 1 and Changes Across Stage 2 Surgery in Single Ventricle Patients. IEEE Trans Biomed Eng. 2018 Nov;65(11):2450–8. doi:10.1109/TBME.2018.2797999. PMID: 29993472.

45. Regazzoni F, Salvador M, Africa PC, Fedele M, Dede L, Quarteroni A. A cardiac electromechanical model coupled with a lumped-parameter model for closed-loop blood circulation. J Comput Phys. 2022 May;457:111083. doi:10.1016/j.jcp.2022.111083.

46. Hoit BD. Anatomy and Physiology of the Pericardium. Cardiol Clin. 2017 Nov;35(4):481–90. doi:10.1016/j.ccl.2017.07.002. PMID: 29025540.

47. Sun Y, Beshara M, Lucariello RJ, Chiaramida SA. A comprehensive model for right-left heart interaction under the influence of pericardium and baroreflex. Am J Physiol. 1997 Mar;272(3 Pt 2):H1499–515. doi:10.1152/ajpheart.1997.272.3.H1499. PMID: 9087629.

48. Nygren A, Fiset C, Firek L, Clark JW, Lindblad DS, Clark RB, et al. Mathematical model of an adult human atrial cell: the role of K+ currents in repolarization. Circ Res. 1998 Jan;82(1):63–81. doi:10.1161/01.res.82.1.63. PMID: 9440706.

49. West JB, Luks A. West’s respiratory physiology: the essentials. Eleventh edition ed. Philadelphia: Wolters Kluwer; 2021.

50. Stickland MK, Lindinger MI, Olfert IM, Heigenhauser GJF, Hopkins SR. Pulmonary gas exchange and acid-base balance during exercise. Compr Physiol. 2013 Apr;3(2):693–739. doi:10.1002/cphy.c110048. PMID: 23720327.

51. Boron WF, Boulpaep EL, editors. Medical physiology. Third edition ed. Philadelphia, PA: Elsevier; 2017.

52. Marshall JM. Peripheral chemoreceptors and cardiovascular regulation. Physiol Rev. 1994 Jul;74(3):543–94. doi:10.1152/physrev.1994.74.3.543. PMID: 8036247.

53. Vadhan J, Tadi P. Physiology, Herring Breuer Reflex. In: StatPearls. Treasure Island (FL): StatPearls Publishing; 2026. PMID: 31869189.

54. St Croix CM, Satoh M, Morgan BJ, Skatrud JB, Dempsey JA. Role of respiratory motor output in within-breath modulation of muscle sympathetic nerve activity in humans. Circ Res. 1999 Sep;85(5):457–69. doi:10.1161/01.res.85.5.457. PMID: 10473675.

55. Raven PB, Fadel PJ, Ogoh S. Arterial baroreflex resetting during exercise: a current perspective. Exp Physiol. 2006 Jan;91(1):37–49. doi:10.1113/expphysiol.2005.032250. PMID: 16210446.

56. Fincham WF, Tehrani FT. A mathematical model of the human respiratory system. J Biomed Eng. 1983 Apr;5(2):125–33. doi:10.1016/0141-5425(83)90030-4. PMID: 6406766.

57. Nelder JA, Mead R. A simplex method for function minimization. The Computer Journal. 1965;7(4):308–13. doi:10.1093/comjnl/7.4.308.

58. Lam SK, Pitrou A, Seibert S. Numba: a LLVM-based Python JIT compiler. In: Proceedings of the Second Workshop on the LLVM Compiler Infrastructure in HPC. LLVM ‘15. New York, NY, USA: Association for Computing Machinery; 2015. p. 1–6. doi:10.1145/2833157.2833162.

59. Verbraecken J, Van de Heyning P, De Backer W, Van Gaal L. Body surface area in normalweight, overweight, and obese adults. A comparison study. Metabolism. 2006 Apr;55(4):515–24. doi:10.1016/j.metabol.2005.11.004. PMID: 16546483.

60. Grewal J, McCully RB, Kane GC, Lam C, Pellikka PA. Left Ventricular Function and Exercise Capacity. JAMA. 2009 Jan;301(3):286–94. doi:10.1001/jama.2008.1022. PMID: 19155455.

61. Miyai N, Arita M, Miyashita K, Morioka I, Shiraishi T, Nishio I. Blood Pressure Response to Heart Rate During Exercise Test and Risk of Future Hypertension. Hypertension. 2002 Mar;39(3):761–6. doi:10.1161/hy0302.105777. PMID: 11897759.

62. Schnell F, Claessen G, La Gerche A, Claus P, Bogaert J, Delcroix M, et al. Atrial volume and function during exercise in health and disease. J Cardiovasc Magn Reson. 2017 Dec;19(1):104. doi:10.1186/s12968-017-0416-9. PMID: 29254488.

63. Haleem SM, Sharma T, Chaudhari SS. Right Heart Catheterization. In: StatPearls. Treasure Island (FL): StatPearls Publishing; 2026. PMID: 32491336.

64. Armstrong DWJ, Matangi MF. Estimated right ventricular systolic pressure during exercise stress echocardiography in patients with suspected coronary artery disease. Can J Cardiol. 2010 Feb;26(2):e45–9. doi:10.1016/S0828-282X(10)70006-4. PMID: 20151058.

65. Cornwell WK, Coe G, Ambardekar A, Pal J, Tompkins C, Zipse M, et al. Abstract 13179: New Insights Into Right Ventricular Performance During Exercise Using High-Fidelity Conductance Catheters to Generate Pressure Volume Loops. Circulation. 2018 Nov;138(Suppl 1):A13179–9. doi:10.1161/circ.138.suppl_1.13179.

66. Karunanithi Z, Andersen MJ, Mellemkjær S, Alstrup M, Waziri F, Skibsted Clemmensen T, et al. Elevated Left and Right Atrial Pressures Long-Term After Atrial Septal Defect Correction: An Invasive Exercise Hemodynamic Study. J Am Heart Assoc. 2021 Jul;10(14):e020692. doi:10.1161/JAHA.120.020692. PMID: 34259012.

67. Namana V, Gupta SS, Sabharwal N, Hollander G. Clinical significance of atrial kick. QJM. 2018 Aug;111(8):569–70. doi:10.1093/qjmed/hcy088. PMID: 29750254.

68. Gabrielli L, Bijnens BH, Brambila C, Duchateau N, Marin J, Sitges-Serra I, et al. Differential atrial performance at rest and exercise in athletes: Potential trigger for developing atrial dysfunction? Scand J Med Sci Sports. 2016 Dec;26(12):1444–54. doi:10.1111/sms.12610. PMID: 26752626.

69. Wright SP, Goodman JM, Sasson Z, Granton JT, Mak S. Left atrial reservoir pressure-volume relations during exercise in healthy older adults. J Appl Physiol (1985). 2024 Apr;136(4):901–7. doi:10.1152/japplphysiol.00905.2023. PMID: 38420677.

70. Sharman JE, Qasem AM, Hanekom L, Gill DS, Lim R, Marwick TH. Radial pressure waveform dP/dt max is a poor indicator of left ventricular systolic function. Eur J Clin Invest. 2007;37(4):276–81. doi:10.1111/j.1365-2362.2007.01784.x. PMID: 17373963.

71. Cornwell WK, Tran T, Cerbin L, Coe G, Muralidhar A, Hunter K, et al. New insights into resting and exertional right ventricular performance in the healthy heart through real-time pressure-volume analysis. J Physiol. 2020;598(13):2575–87. doi:10.1113/JP279759. PMID: 32347547.

72. Peel JK, Funk DJ, Slinger P, Srinathan S, Kidane B. Tidal volume during 1-lung ventilation: A systematic review and meta-analysis. J Thorac Cardiovasc Surg. 2022 Apr;163(4):1573-85.e1. doi:10.1016/j.jtcvs.2020.12.054. PMID: 33518385.

73. Orton CM, Symons HE, Moseley B, Archer J, Watson NA, Philip KEJ, et al. A comparison of respiratory particle emission rates at rest and while speaking or exercising. Commun Med (Lond). 2022 Apr;2(1):44. doi:10.1038/s43856-022-00103-w. PMID: 35603287.

74. Wheatley CM, Snyder EM, Johnson BD, Olson TP. Sex differences in cardiovascular function during submaximal exercise in humans. Springerplus. 2014 Aug;3(1):445. doi:10.1186/2193-1801-3-445. PMID: 25191635.

75. Stoffel MA, Li BM, Westerling K, Arana S, Balmus M, Daub E, et al. AutoEmulate: A Python package for semi-automated emulation. J Open Source Softw. 2025 Mar;10(107):7626. doi:10.21105/joss.07626.

76. Sklar A. Random variables, joint distribution functions, and copulas. Kybernetika (Prague). 1973;9(6):449–60. Available from: https://eudml.org/doc/28992.

77. Zhang L, Singh VP. Copulas and their applications in water resources engineering. Cambridge New York, NY: Cambridge University Press; 2019.

78. Hoffman MD, Gelman A. The No-U-Turn Sampler: Adaptively Setting Path Lengths in Hamiltonian Monte Carlo. J Mach Learn Res. 2014;15(47):1593–623. Available from: http://jmlr.org/papers/v15/hoffman14a.html.

79. Gelman A, Rubin DB. Inference from Iterative Simulation Using Multiple Sequences. Stat Sci. 1992 Nov;7(4):457–72. doi:10.1214/ss/1177011136.

80. Geyer CJ. Practical Markov Chain Monte Carlo. Stat Sci. 1992 Nov;7(4):473–83. doi:10.1214/ss/1177011137.

81. Kucherenko S, Klymenko OV, Shah N. Sobol’ indices for problems defined in non-rectangular domains. Reliab Eng Syst Saf. 2017 Nov;167:218–31. doi:10.1016/j.ress.2017.06.001.

82. Hilhorst PLJ, Quicken S, van de Vosse FN, Huberts W. Efficient sensitivity analysis for biomechanical models with correlated inputs. Int J Numer Method Biomed Eng. 2024;40(2):e3797. doi:10.1002/cnm.3797. PMID: 38116742.

83. Hilhorst P, van de Wouw B, Zajac K, van ‘t Veer M, Tonino P, van de Vosse F, et al. Sensitivity analysis for exploring the variability and parameter landscape in virtual patient cohorts of multivessel coronary artery disease. Philos Trans A Math Phys Eng Sci. 2025 Apr;383(2293):20240230. doi:10.1098/rsta.2024.0230. PMID: 40172563.

84. Vehtari A, Gelman A, Simpson D, Carpenter B, Bürkner PC. Rank-Normalization, Folding, and Localization: An Improved R^^^ for Assessing Convergence of MCMC (with Discussion). Bayesian Anal. 2021 Jun;16(2):667–718. doi:10.1214/20-BA1221.

85. Sato K, Ogoh S, Hirasawa A, Oue A, Sadamoto T. The distribution of blood flow in the carotid and vertebral arteries during dynamic exercise in humans. J Physiol. 2011 Jun;589(Pt 11):2847–56. doi:10.1113/jphysiol.2010.204461. PMID: 21486813.

86. Sheriff DD. Baroreflex resetting during exercise: mechanisms and meaning. Am J Physiol Heart Circ Physiol. 2006 Apr;290(4):H1406–7. doi:10.1152/ajpheart.01275.2005. PMID: 16537789.

87. White DW, Raven PB. Autonomic neural control of heart rate during dynamic exercise: revisited. J Physiol. 2014;592(12):2491–500. doi:10.1113/jphysiol.2014.271858. PMID: 24756637.

88. Perko MJ, Nielsen HB, Skak C, Clemmesen JO, Schroeder TV, Secher NH. Mesenteric, coeliac and splanchnic blood flow in humans during exercise. J Physiol. 1998;513(3):907–13. doi:10.1111/j.1469-7793.1998.907ba.x. PMID: 9824727.

89. Forster HV, Haouzi P, Dempsey JA. Control of breathing during exercise. Compr Physiol. 2012 Jan;2(1):743–77. doi:10.1002/cphy.c100045. PMID: 23728984.

90. Ward SA, Whipp BJ, Koyal S, Wasserman K. Influence of body CO2 stores on ventilatory dynamics during exercise. J Appl Physiol Respir Environ Exerc Physiol. 1983 Sep;55(3):742–9. doi:10.1152/jappl.1983.55.3.742. PMID: 6415010.

91. Chuang ML, Ting H, Otsuka T, Sun XG, Chiu FY, Beaver WL, et al. Aerobically generated CO(2) stored during early exercise. J Appl Physiol (1985). 1999 Sep;87(3):1048–58. doi:10.1152/jappl.1999.87.3.1048. PMID: 10484576.

92. Proctor DN, Beck KC, Shen PH, Eickhoff TJ, Halliwill JR, Joyner MJ. Influence of age and gender on cardiac output-Vo 2 relationships during submaximal cycle ergometry. J Appl Physiol (1985). 1998 Feb;84(2):599–605. doi:10.1152/jappl.1998.84.2.599. PMID: 9475871.

93. Schweitzer R, de Marvao A, Shah M, Inglese P, Kellman P, Berry A, et al. Establishing Cardiac MRI Reference Ranges Stratified by Sex and Age for Cardiovascular Function during Exercise. Radiol Cardiothorac Imaging. 2025 Jun;7(3):e240175. doi:10.1148/ryct.240175. PMID: 40471075.

94. Wolsk E, Bakkestrøm R, Thomsen JH, Balling L, Andersen MJ, Dahl JS, et al. The Influence of Age on Hemodynamic Parameters During Rest and Exercise in Healthy Individuals. JACC Heart Fail. 2017 May;5(5):337–46. doi:10.1016/j.jchf.2016.10.012. PMID: 28017352.

95. Beck KC, Johnson BD, Olson TP, Wilson TA. Ventilation-perfusion distribution in normal subjects. J Appl Physiol (1985). 2012 Sep;113(6):872–7. doi:10.1152/japplphysiol.00163.2012. PMID: 22773767.

96. Schwartz JC, Snyder EM, Olson TP, Johnson BD, Wheatley-Guy CM. Alveolar to arterial gas exchange during constant-load exercise in healthy active men and women. J Sports Sci. 2021 May;39(9):961–8. doi:10.1080/02640414.2020.1851927. PMID: 33242298.

97. Radtke T, Crook S, Kaltsakas G, Louvaris Z, Berton D, Urquhart DS, et al. ERS statement on standardisation of cardiopulmonary exercise testing in chronic lung diseases. Eur Respir Rev. 2019 Dec;28(154). doi:10.1183/16000617.0101-2018. PMID: 31852745.

98. Qian S, Ugurlu D, Fairweather E, Toso LD, Deng Y, Strocchi M, et al. Developing cardiac digital twin populations powered by machine learning provides electrophysiological insights in conduction and repolarization. Nat Cardiovasc Res. 2025 May;4(5):624–36. doi:10.1038/s44161-025-00650-0. PMID: 40379795.

99. Saltelli A, Annoni P. How to avoid a perfunctory sensitivity analysis. Environ Model Softw. 2010 Dec;25(12):1508–17. doi:10.1016/j.envsoft.2010.04.012.

100. Pianosi F, Beven K, Freer J, Hall JW, Rougier J, Stephenson DB, et al. Sensitivity analysis of environmental models: A systematic review with practical workflow. Environ Model Softw. 2016 May;79:214–32. doi:10.1016/j.envsoft.2016.02.008.

101. Arts T, Delhaas T, Bovendeerd P, Verbeek X, Prinzen FW. Adaptation to mechanical load determines shape and properties of heart and circulation: the CircAdapt model. Am J Physiol Heart Circ Physiol. 2005 Apr;288(4):H1943–54. doi:10.1152/ajpheart.00444.2004. PMID: 15550528.

102. Williams L, Frenneaux M. Diastolic ventricular interaction: from physiology to clinical practice. Nat Clin Pract Cardiovasc Med. 2006 Jul;3(7):368–76. doi:10.1038/ncpcardio0584. PMID: 16810172.

103. Leenders GE, Lumens J, Cramer MJ, De Boeck BWL, Doevendans PA, Delhaas T, et al. Septal deformation patterns delineate mechanical dyssynchrony and regional differences in contractility: analysis of patient data using a computer model. Circ Heart Fail. 2012 Jan;5(1):87–96. doi:10.1161/CIRCHEARTFAILURE.111.962704. PMID: 21980078.

104. Angus SA, Taylor JL, Mann LM, Williams AM, Stöhr EJ, Au JS, et al. Attenuating intrathoracic pressure swings decreases cardiac output at different intensities of exercise. J Physiol. 2023 Nov;601(21):4807–21. doi:10.1113/JP285101. PMID: 37772933.

105. Menuet C, Ben-Tal A, Linossier A, Allen AM, Machado BH, Moraes DJA, et al. Redefining respiratory sinus arrhythmia as respiratory heart rate variability: an international Expert Recommendation for terminological clarity. Nat Rev Cardiol. 2025 Dec;22(12):978–84. doi:10.1038/s41569-025-01160-z. PMID: 40328963.

106. van Heusden K, Gisolf J, Stok WJ, Dijkstra S, Karemaker JM. Mathematical modeling of gravitational effects on the circulation: importance of the time course of venous pooling and blood volume changes in the lungs. Am J Physiol Heart Circ Physiol. 2006 Nov;291(5):H2152–65. doi:10.1152/ajpheart.01268.2004. PMID: 16632542.

107. Schiavazzi DE, Arbia G, Baker C, Hlavacek AM, Hsia TY, Marsden AL, et al. Uncertainty quantification in virtual surgery hemodynamics predictions for single ventricle palliation. Int J Numer Method Biomed Eng. 2016 Mar;32(3):e02737. doi:10.1002/cnm.2737. PMID: 26217878.

108. Colebank MJ, Umar Qureshi M, Olufsen MS. Sensitivity analysis and uncertainty quantification of 1-D models of pulmonary hemodynamics in mice under control and hypertensive conditions. Int J Numer Method Biomed Eng. 2021 Nov;37(11):e3242. doi:10.1002/cnm.3242. PMID: 31355521.

109. Argus F, Zhao D, Babarenda Gamage TP, Nash MP, Maso Talou GD. Automated model calibration with parallel MCMC: Applications for a cardiovascular system model. Front Physiol. 2022 Nov;13. doi:10.3389/fphys.2022.1018134. PMID: 36439250.

110. Hu Z, Wang H, Zhou Q. A MCMC method based on surrogate model and Gaussian process parameterization for infinite Bayesian PDE inversion. J Comput Phys. 2024 Jun;507:112970. doi:10.1016/j.jcp.2024.112970.

111. Paun LM, Colebank MJ, Olufsen MS, Hill NA, Husmeier D. Assessing model mismatch and model selection in a Bayesian uncertainty quantification analysis of a fluid-dynamics model of pulmonary blood circulation. J R Soc Interface. 2020 Dec;17(173):20200886. doi:10.1098/rsif.2020.0886. PMID: 33353505.

112. Sürer Plumlee M, Wild SM. Sequential Bayesian Experimental Design for Calibration of Expensive Simulation Models. Technometrics. 2024 Apr;66(2):157–71. doi:10.1080/00401706.2023.2246157.

113. Miura T, Miyazaki S, Guth BD, Kambayashi M, Ross J. Influence of the force-frequency relation on left ventricular function during exercise in conscious dogs. Circulation. 1992 Aug;86(2):563–71. doi:10.1161/01.CIR.86.2.563. PMID: 1638722.

114. Marquis AD, Arnold A, Dean-Bernhoft C, Carlson BE, Olufsen MS. Practical identifiability and uncertainty quantification of a pulsatile cardiovascular model. Math Biosci. 2018 Oct;304:9–24. doi:10.1016/j.mbs.2018.07.001. PMID: 30017910.

115. Sarmiento CA, Serna LY, Hernández AM, Mañanas MA. A Novel Strategy to Fit and Validate Physiological Models: A Case Study of a Cardiorespiratory Model for Simulation of Incremental Aerobic Exercise. Diagnostics (Basel). 2023 Jan;13(5):908. doi:10.3390/diagnostics13050908. PMID: 36900052.

116. Kung E, Perry JC, Davis C, Migliavacca F, Pennati G, Giardini A, et al. Computational modeling of pathophysiologic responses to exercise in Fontan patients. Ann Biomed Eng. 2015 Jun;43(6):1335–47. doi:10.1007/s10439-014-1131-4. PMID: 25260878.

